# High-throughput *in silico* screen uncovers key regulators of 3D genome architecture

**DOI:** 10.64898/2025.12.09.693120

**Authors:** Jiangshan Bai, Qingji Lyu, Jimin Tan, Robbyn Issner, Bailey Tischer, Hanqing Liu, Viraat Goel, Xinyu Ling, Bradley E. Bernstein, Aristotelis Tsirigos, Anders S. Hansen, Bo Xia

## Abstract

The vertebrate genome is spatially organized into topologically associating domains (TADs), primarily via cohesin-mediated loop extrusion which typically halts at convergent CTCF binding sites to establish domain boundaries. However, despite the essential roles of CTCF and cohesin in establishing TADs, a long-standing paradox persists: CTCF and cohesin binding sites dramatically outnumber observed TAD boundaries, suggesting the existence of undiscovered architectural factors. To identify such missing factors, we conducted high-resolution *in silico* screens using C.Origami, a multi-modal AI model for predicting chromatin interactions. Remarkably, we identified ZNF654 and JMJD6 as novel factors uniquely defining TAD boundaries. Experimental validation confirmed that ZNF654, an uncharacterized vertebrate-specific zinc-finger protein, interacts with CTCF to form an architectural protein complex that demarcates chromatin domains. Genetic knockout of *ZNF654* weakens TAD boundary strength without influencing other CTCF or cohesin binding sites. JMJD6, a deeply conserved jmjC-family dioxygenase, marks the anchors of the strongest chromatin stripes at both TAD boundaries and enhancer-promoter sites, while deleting *JMJD6* weakens or diminishes such interaction signature. These results revealed the long-sought factors that uniquely mark TAD boundary and chromatin interaction anchors which, together with CTCF and cohesin, demarcate chromatin domains during 3D genome organization. Last, the evolutionary trajectory of ZNF654 and JMJD6 offers key insight into the evolutionary origins of 3D genome organization across metazoan species.

## INTRODUCTION

Proper regulation of gene expression requires the genome to fold into organized three-dimensional conformations, bringing regulatory elements and their target genes into precise spatial proximity^1,2^. High-resolution chromosome conformation capture assays such as Hi-C and Micro-C have revealed a rich hierarchy of genome conformational structures – compartments, topologically associating domains (TADs), loops, and stripes – that scaffold long-range gene regulatory circuits^3–8^. The loop extrusion model, in which cohesin complex extrudes chromatin until impeded by convergent CTCF-occupied sites, provides a key mechanistic foundation for three-dimensional (3D) genome architecture at the scale of TADs and loops^9–14^.

Yet several long-standing inconsistencies indicate that the current loop extrusion model is incomplete. First, CTCF and cohesin occupy far more sites (∼50,000) than the observed number of TAD boundaries (∼10,000) in the human genome. Thus, most CTCF-and cohesin-occupied sites do not form strong boundaries^4,5,15–18^. Second, acute CTCF depletion weakens TAD boundaries but leaves the majority of domain architecture intact. In contrast, cohesin depletion disrupts most TAD structures and cohesin-associated loops and stripes, yet still spares many microcompartment-like contacts, transcription-linked stripes, and global gene expression^19–22^. These observations highlight a critical unresolved question: what additional factors could uniquely specify TAD boundary sites and stabilize transcription-associated chromatin contacts?

Attempts to identify such regulators through biochemical assays, genetic or optical screen approaches have proposed candidates such as ZFP143, MAZ, YY1, PATZ1, and GSK3A as putative regulators of chromatin interactions^23–27^. However, the binding profiles of these factors do not enrich at the observed TAD boundaries, suggesting that their functions do not extend to shape global 3D genome conformation. Notably, recent studies confirmed that ZNF143, a mischaracterized looping factor, functions as a transcription factor binding primarily at promoters but not TAD boundaries^28,29^. Therefore, the general molecular mechanisms by which TAD boundaries are defined and established remains largely unresolved.

Here, we leveraged a deep-learning-based *in silico* genetic screen approach that precisely predicts how kilobase-scale perturbations reshape 3D genome conformation^30^. Through interpreting the *in silico* screen-identified key genomic elements to infer novel 3D genome regulators, we discovered two previously uncharacterized regulators – ZNF654, a vertebrate-specific zinc-finger protein that interacts with CTCF to uniquely reinforce TAD boundaries, and JMJD6, a deeply conserved jmjC-family dioxygenase that stabilizes both boundary anchors and enhancer-promoter stripes. Perturbing ZNF654 weakens TAD boundary insulation and cohesin-mediated loop strength, while JMJD6 loss disrupts chromatin loops and stripes at both TAD boundary and promoter-enhancer sites. Together, these findings revealed the long-sought missing components of the molecular machinery that uniquely define TAD boundaries and chromatin stripes. To our knowledge, this is the first time that a deep learning-based *in silico* experimentation and screen approach leads to the discovery of critical regulators in genome biology, highlighting a powerful AI-enabled framework that complements experimental approaches to accelerate biological discoveries.

## RESULTS

### High-resolution *in silico* screen identifies ZNF654 and JMJD6 as top candidate regulators of 3D genome conformation

To uncover previously unrecognized regulators of 3D genome organization, we carried out a high-resolution *in silico* screen leveraging our multi-modal deep neural network, C.Origami^30^. We first performed chromosome-wide saturated *in silico* deletions to systematically quantify how fine-scale genomic elements contribute to 3D genome conformation (Fig. 1a, Methods). This *in silico* screen strategy enabled precise identification of kilobase-scale genomic loci whose perturbation strongly alters boundary insulation or inter-TAD interactions – a capability that would not be feasible through conventional experimental approaches.

**Figure 1.**
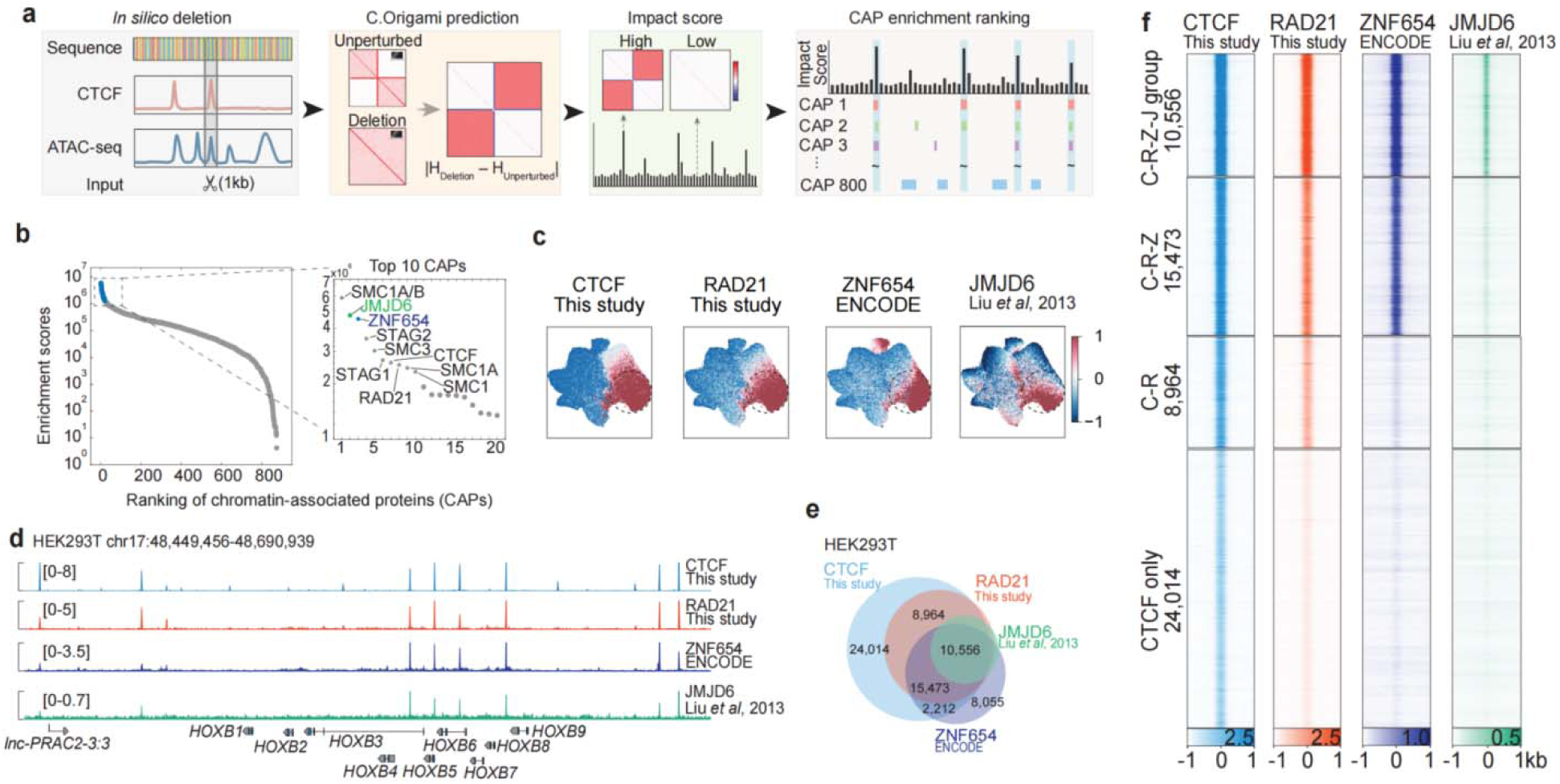
*In silico* screen identified ZNF654 and JMJD6 as top candidates for 3D genome regulation. **a**, Schematic of the high-throughput *in silico* deletion screen workflow using C.Origami. **b**, Ranking of chromatin proteins by their enriched binding at structurally high-impact regions, with the top 10 factors highlighted. **c**, UMAP visualization of ChIP-seq signal intensities for CTCF, RAD21, ZNF654, and JMJD6 across ∼100,000 key CREs. **d**, Genomic track view at the *HOXB* gene cluster locus in HEK293T cells, showing ChIP-seq signals for CTCF, RAD21, ZNF654, and JMJD6. **e**, Venn diagram showing overlap of CTCF, RAD21, ZNF654, and JMJD6 occupied sites. **f**, Heatmaps of ChIP-seq signal centered on CTCF sites grouped by co-occupancy with RAD21, ZNF654, and JMJD6.

Next, we integrated binding profiles of all chromatin-associated proteins from the ReMap database^31^ and ranked factors by their enrichment of binding at high structural impact regions (Fig. 1a). As expected, core architectural protein CTCF and canonical loop-extrusion components in the cohesin complexes, including RAD21, STAG1, STAG2, SMC1, and SMC3 – were among the top-ranked factors (Fig. 1b). Surprisingly, two previously uncharacterized proteins also emerged as top candidates: JMJD6, a highly conserved jmjC-family dioxygenase with debating enzymatic functions ^32–37^; and ZNF654, an uncharacterized, vertebrate-specific C2H2 zinc-finger protein.

To systematically characterize the genomic binding patterns of key chromatin factors, we integrated ChIP-seq profiles for CTCF, RAD21, ZNF654, JMJD6, H3K27ac, H3K4me3, Pol II, and ATAC-seq into a unified set of ∼100,000 key *cis*-regulatory elements (CREs, Methods). We quantified the intensities of these chromatin features at each element and visualized them using UMAP embedding, enabling a direct comparison of feature co-occupancy patterns across these key CREs (Fig. 1c and S1). This analysis revealed a prominent cluster of CREs uniquely enriched for JMJD6 and ZNF654, occupying a subset of CTCF and cohesin binding sites (Fig. 1c and S1a). Notably, ZNF143, YY1, MAZ, and PATZ1 ^23–26^, tend to localize near, but not extensively overlap with the CTCF-and cohesin-bound CREs (Fig. S1b), reflecting their more selective or context-dependent roles on chromatin interactions.

Strikingly, we found that JMJD6 and ZNF654 exhibit nearly exclusive co-occupancy with CTCF and cohesin across the genome (Fig. 1c-d). To rigorously validate these observed co-occupancy patterns, we performed a series of independent analyses and experiments. First, we confirmed that JMJD6 consistently co-localizes with CTCF and cohesin across different human cell types, and such co-localization pattern is conserved in mouse cells where the ChIP-seq data was collected using a different antibody (Fig. S2). Second, to further validate these observations beyond public datasets, we generated endogenous epitope-tagged cell lines for both factors and screened additional antibodies recognizing distinct epitopes. ChIP-seq using anti-FLAG and a newly validated antibody recognizing N-terminal peptides of JMJD6 recapitulated its genome-wide binding sites in both HEK293T and HCT116 cells (Fig. S3). These independent datasets also revealed another group of JMJD6 binding at enhancer-promoter sites (that will be discussed later). Similarly, anti-HA ChIP-seq in an endogenously HA-tagged ZNF654 line in HCT116 cells revealed the consistent co-occupancy of ZNF654 with CTCF and cohesin (Fig. S4), same as that observed in HEK293T cells (Fig. 1c-d). Collectively, these results confirm that JMJD6 and ZNF654 bind at a distinct subset of CTCF-occupied sites.

### ZNF654 uniquely defines a boundary-forming subclass of CTCF sites

To determine whether ZNF654 and JMJD6 mark the subset of CTCF sites that form bona fide TAD boundaries, we first classified CTCF-occupied sites into four categories based on factor co-occupancy: regions bound by all four factors (CTCF, RAD21, ZNF654, and JMJD6; the ‘C–R–Z–J’ group), by three factors (except JMJD6, ‘C–R–Z’ group), by CTCF and RAD21 only (‘C–R’ group), or by CTCF alone (‘CTCF-only’ group) (Fig. 1f). Notably, the ∼10,000 C–R–Z–J sites approximate the number of TAD boundaries in the human genome, representing a selectively enriched subset of the ∼50,000 CTCF-occupied sites genome-wide (Fig. 1f).

We next aligned and piled deeply sequenced Micro-C contact maps to each CTCF-occupied category. In HEK293 cells, C–R–Z–J sites exhibited the strongest insulation signals and followed by the C–R–Z sites (Fig. 2a-b), both marking robust TAD boundaries, whereas C–R and CTCF-only sites do not show insulating boundary features (Fig. 2a-b). Analysis of loop-anchored interactions further revealed that both C–R–Z–J and C–R–Z sites exhibited prominent loop extrusion signatures, characterized by focal contacts flanked by broad stripes extending from anchors (Fig. 2c). Cohesin loop strength progressively diminished with the loss of JMJD6 or ZNF654 binding across the remaining groups, indicating that these two factors are key predictors of extrusion competence (Fig. 2c). Analyses using independent datasets in HCT116 cells reproduced the same hierarchy, with C–R–Z–J sites marking the strongest boundaries followed by C–R–Z sites (Fig. S5). These results suggest that ZNF654 binding defines the TAD boundary, while additional binding by JMJD6 further strengthens the boundary.

**Figure 2.**
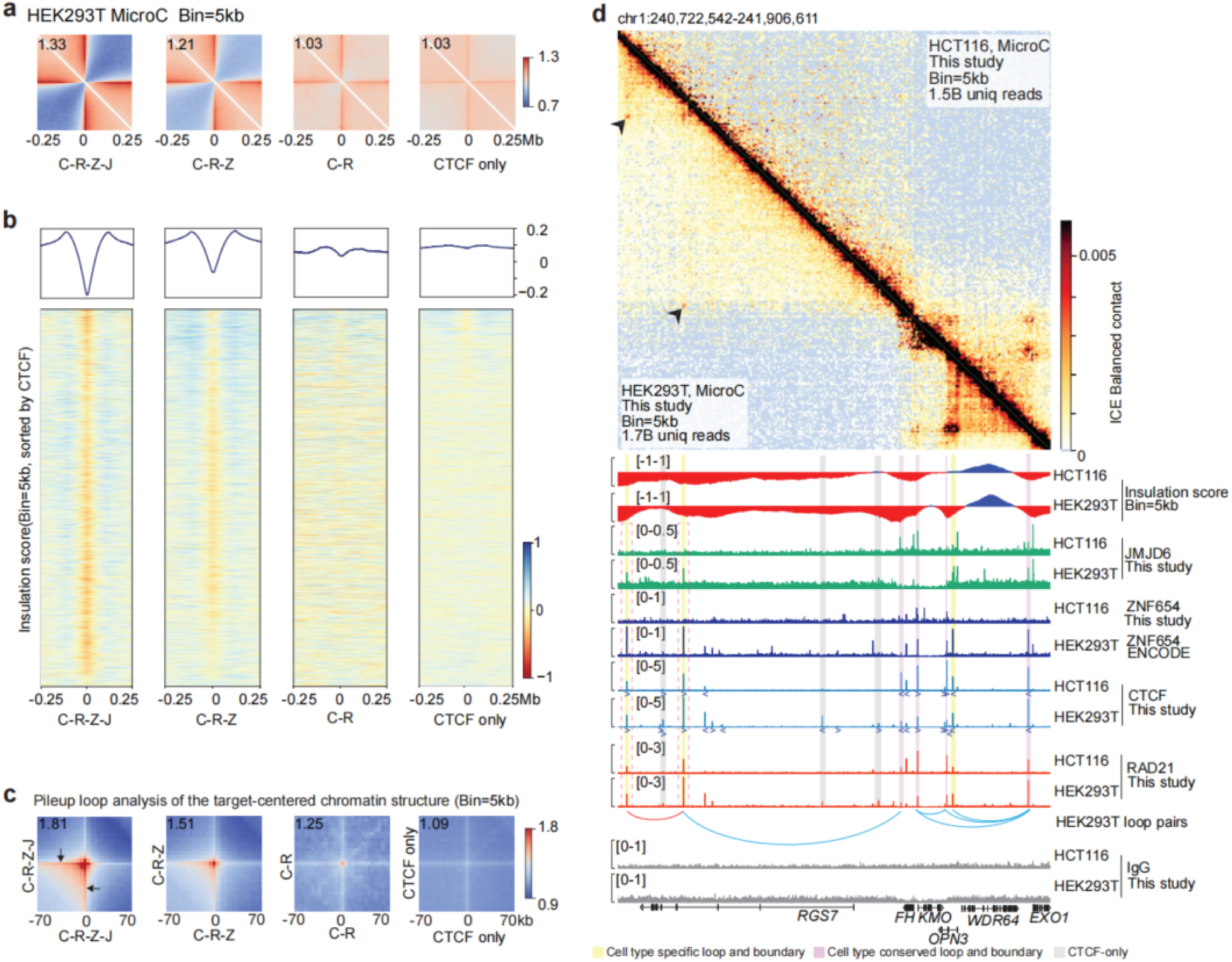
ZNF654 and JMJD6 mark the strong TAD boundaries and loop anchors **a**, Micro-C pile-up visualization of chromatin interaction intensities flanking the loci belonging to the four CTCF-occupied groups. For genomic loci in each group, the Micro-C contact map is centered at the factor binding site, and visualized at 5Kb/pixel resolution. **b**, Heatmap visualization of insulation scores at each genomic locus (5-kb resolution). Loci in each category was sorted by CTCF strength. Average Insulation score profile was displayed on top of each heatmap. **c**, Aggregate peak analysis (APA) visualization of chromatin loops centered on peak pairs within each binding category. **d**, Aligned visualization of Micro-C contact maps and ChIP-seq tracks at a representative genomic region (chr1:240.7–241.9 Mb) in HEK293T and HCT116 cells. Genomic loci marking cell-type-specific and conserved structural sites are highlighted in yellow and purple, respectively. Other CTCF binding sites are marked in grey.

Importantly, ZNF654 binding defines cell-type-specific TAD boundaries. Parallel alignment of ChIP-seq and Micro-C data across HEK293, HCT116, and HeLa cells confirmed that ZNF654-bound sites consistently correspond to the most prominent TAD boundaries and loop anchors, and cell-type-specific TAD boundaries are distinguished by cell-type-specific ZNF654-binding but not CTCF or cohesin binding (Fig. 2d and S6). For example, HEK293-specific ZNF654 binding sites corresponded to HEK293-specific boundaries and loops that were absent in HCT116, despite comparable CTCF and RAD21 occupancy at these loci (Fig. 2d and S6). Conversely, regions with conserved ZNF654 binding exhibited shared TAD boundary formation across cell types, demonstrating that the presence or absence of ZNF654 – but not the differences in CTCF or cohesin binding – underlies cell-type-specific TAD boundary formation (Fig. 2d and S6).

### Classical CTCF features do not explain boundary strength

We next asked whether classical sequence or chromatin features could account for the differential insulation strength observed across the four CTCF-occupied loci groups. First, motif analyses across the four groups revealed almost identical CTCF sequence motifs (Fig. S7a). Similarly, motif scores showed no categorial difference among groups – strong boundaries within the C–R–Z–J and C–R–Z groups have a similarly broad spectrum of CTCF motif scores ranging from strong to weak in comparison to those in the non-boundary C–R or CTCF-only groups (Fig. S7b). These results suggest that CTCF motif strength alone do not confer strong TAD boundary features ^38,39^.

Recent work proposed that local CTCF clustering – multiple consecutive CTCF peaks within a small genomic locus – may strengthen TAD boundaries^38,40,41^. We noticed that these observations typically derive from analysis of boundary calling in low-resolution Hi-C maps (Fig. S7c). To test whether C–R–Z–J and C–R–Z co-occupied sites – the TAD boundaries – have higher density of CTCF binding, we next quantified the number of nearby CTCF peaks for each of the CTCF-occupied sites (Methods). We found CTCF density across groups was indistinguishable within genomic windows ranging from ±1 kb to ±100 kb (Fig. S7d), indicating that clustering does not underlie the differential insulation strength observed across CTCF-occupied groups.

Last, we observed a counterintuitive inverse relationship between chromatin accessibility and boundary strength. When stratifying C–R–Z–J and C–R–Z sites by ATAC-seq signal intensity, insulation increased as accessibility decreased (Fig. S7e). Binding intensities of CTCF, RAD21, ZNF654, and JMJD6 were overall stable across accessibility quartiles, suggesting that the TAD boundary formation involving ZNF654 and JMJD6 is not dictated by the strength of chromatin openness.

Together, these analyses show that CTCF motif strength, motif conservation, CTCF clustering, and chromatin accessibility cannot explain explicitly why only a subset of CTCF-occupied sites forms strong TAD boundaries. Instead, our results indicate that co-occupancy with ZNF654 and JMJD6 is the key predictor of boundary activity.

### ZNF654 strengthens TAD boundaries through interacting with CTCF

ZNF654 uniquely binds at high-insulating subset of CTCF-occupied sites, suggesting a putative function as an architectural protein that halts cohesin-mediated loop extrusion to establish TAD boundaries. To test this hypothesis, we generated ZNF654 knockout (KO) lines in HEK293 and HCT116 cells (Fig. 3a-b and S8). Subsequently, we performed high-resolution Micro-C assay, generating 3D chromatin contact maps in both wild-type (WT) and KO cells with comparable sequencing depth and data quality (Fig. S8c-d, Methods).

**Figure 3.**
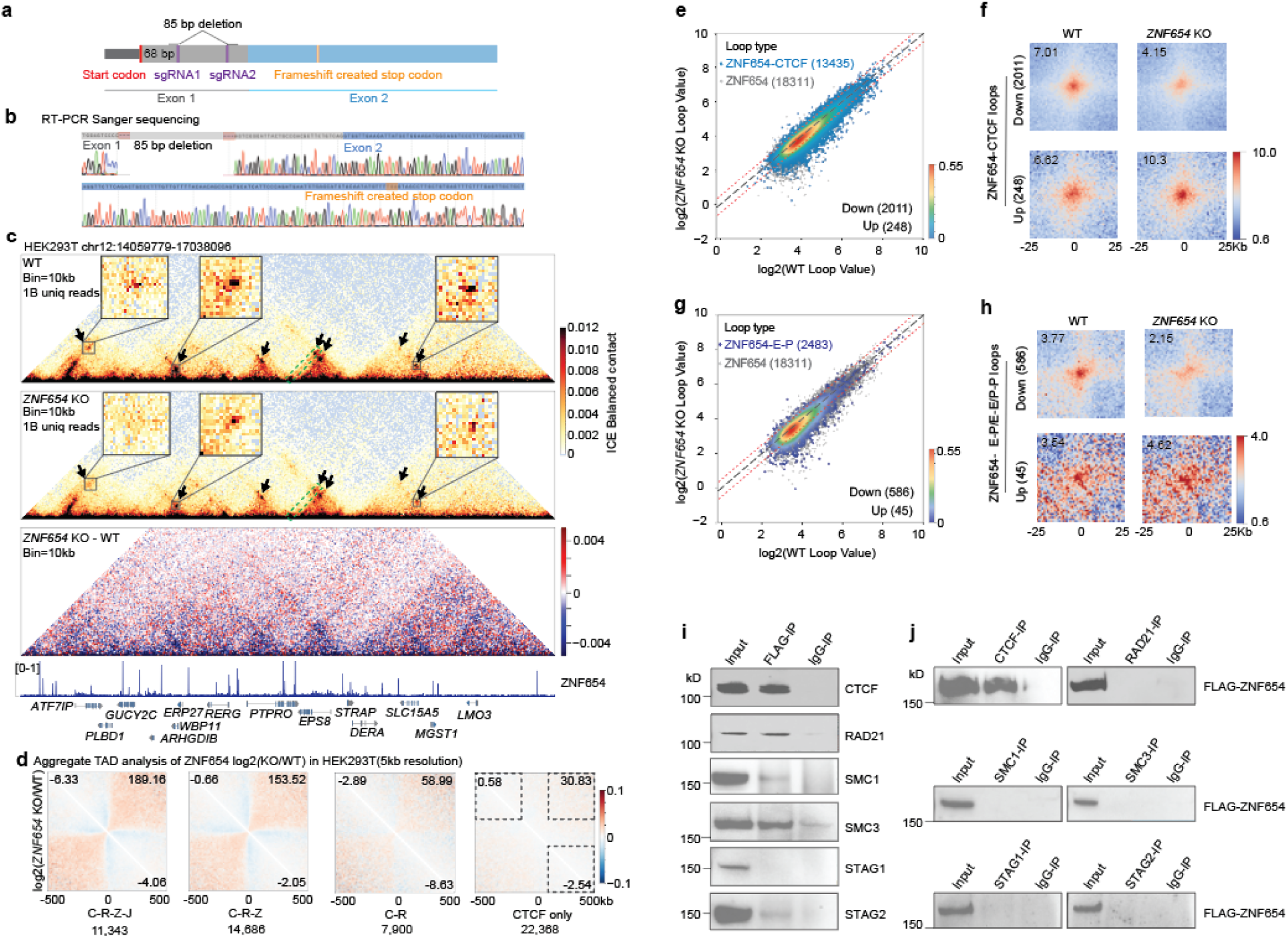
ZNF654 strengthens TAD boundary through interacting with CTCF **a-b**, CRISPR-Cas9 strategy for generating ZNF654 knockout (KO) alleles by dual sgRNA targeting exon 1 of the gene, generating an 85bp deletion that leads to frameshift and early stop codon that disrupt the protein. **c**, Micro-C contact maps in WT (top) and ZNF654-KO (middle) HEK293 cells at 10Kb resolution. Arrows indicate weakened loops in ZNF654-KO cells. Three loops are highlighted and shown at 5 kb resolution in zoomed-in views. Green dashed box indicates a weakened boundary. Difference map (bottom) of chromatin interaction changes (KO relative to WT) shows increased inter-TAD interactions upon knockout of ZNF654. **d**, Pile-up visualization of chromatin interaction changes across the four CTCF-occupied groups, highlighting the dramatic increase of inter-TAD interactions spanning the ZNF654-occupied loci upon knockout of ZNF654. The three corner numbers in each heatmap indicate the total interaction changes in the dash-squared area of each corner. **e**, Scatter plot comparing loop strength between wild-type and ZNF654-KO cells at ZNF654–CTCF co-occupied loop sites, with the number of significantly different loops annotated. **f**, Aggregate peak analysis (APA) visualization of significantly weakened or strengthened ZNF654–CTCF loops. **g-h**, Same as (**e**-**f**) for ZNF654-anchored enhancer–promoter loops. **i**, Anti-FLAG co-immunoprecipitation pull-down followed by CTCF, RAD21, SMC1, SMC3, STAG1 and STAG2 western blot in HCT116 3xFLAG-ZNF654 cells. All proteins are expressed at their endogenous level. **j**, Reciprocal IP pull-down using antibodies against CTCF or cohesin proteins followed by blotting FLAG tag (3xFLAG-ZNF654).

As expected, ZNF654 loss led to widespread weakening of TAD boundary strength and cohesin-mediated loops, represented by increased inter-TAD interactions and concurrent losses in focal contact intensity, respectively (Fig. 3c-h). Genome-wide analyses of boundary strength upon knockout of ZNF654 revealed increased inter-TAD interactions – suggesting decreased boundary strength – only evident in the C–R–Z–J and C–R–Z sites, but not in C–R or CTCF-only sites, confirming a selective loss of insulation at the ZNF654-occupied sites (Fig. 3d and S8e-g). Chromatin loop analysis further showed widespread cohesin-mediated loop weakening at ZNF654 sites upon knockout (Fig. 3e), with 2,011 loops showing significantly reduced intensity versus only 248 increased (Fig. 3f). Similar patterns were observed at ZNF654-bound enhancer–promoter loops despite their relatively low occurrence across the genome (Fig. 3g-h). These phenotypes were consistent in HCT116 cells, establishing a conserved role for ZNF654 in specifying TAD conformation and loops at the subset of CTCF-occupied sites (Fig. S8f-k).

These phenotypic results lead us to hypothesize that ZNF654 works together with CTCF as an architectural protein complex to establish TAD boundary during loop extrusion. To test whether ZNF654 physically interacts with known 3D genome regulators proteins, we performed co-immunoprecipitation (co-IP) in endogenously FLAG-tagged ZNF654 cells (Fig. S8l-m). Endogenously expressed FLAG-ZNF654 protein robustly co-immunoprecipitated with CTCF, RAD21, and SMC3, with weaker recovery of SMC1 and STAG2 (Fig. 3i). Reciprocal co-IP showed that only CTCF efficiently retrieved ZNF654 (Fig. 3j), supporting a direct and stable *in vivo* interaction between ZNF654 and CTCF. These findings suggest that ZNF654 interacts with CTCF to form an architectural protein complex that establish TAD boundaries during loop extrusion.

### JMJD6 marks both TAD boundaries and chromatin stripes

Next, we systematically characterized the function of JMJD6 on 3D genome architecture. Our analysis of public ChIP-seq datasets showed that JMJD6 is strongly enriched at C–R–Z–J sites, the strongest TAD boundaries (Fig. 1-2, S1-2). Using a rigorously validated JMJD6 antibody and endogenous epitope tagging, we confirmed this TAD boundary-binding pattern and, additionally, uncovered another class of JMJD6-bound sites featured by RNA Pol II binding and H3K27ac signals (Fig. 4a-b, Fig. S3e-h, S9a-b). UMAP projection of chromatin signals revealed that the other group of JMJD6 binding is enriched with active chromatin features (H3K27ac, H3K4me3, and Pol II occupancy) and mapped overwhelmingly close to transcription start sites (median distance < 100 bp; Fig. 4b-c, S9c-d). High-resolution Micro-C (e.g. bin size = 500bp) analysis revealed that JMJD6 occupies not only cohesin-mediated extrusion loop anchors but also the anchors of fine-scale promoter-promoter (P–P) and enhancer-promoter (E–P) stripes (Fig. 4a). Cell-type-specific stripes are distinguished by JMJD6 occupation but not by other factors such as CTCF, ZNF654, cohesin or H3K27ac (Fig. 4a and S9e-f). Notably, many JMJD6-marked transcriptional stripes bypassed the cohesin-defined TAD boundary, forming a continuous interaction stripe typically at highly active promoters (Fig. 4a and S9e). These results are consistent with previous observations that transcription-associated chromatin interactions can extend beyond TAD boundaries ^21,42,43^.

**Figure 4.**
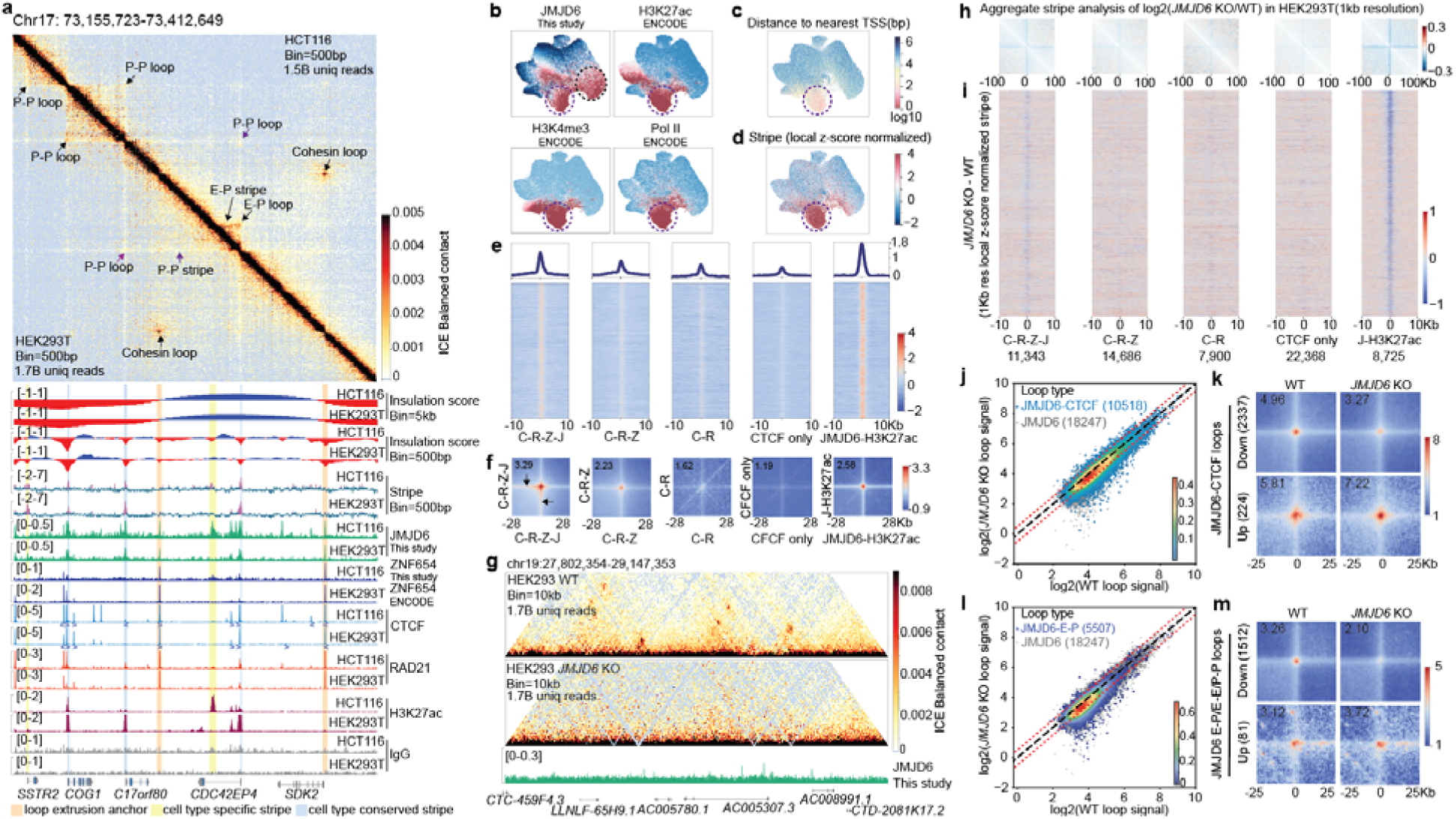
JMJD6 strengthens stripe and loop interactions. **a**, High-resolution Micro-C maps (500bp resolution) comparison between HEK293T and HCT116 cells. Loops and stripes are annotated, with associated ChIP-seq tracks and calculated insulation and stripe scores shown below. Orange highlights mark loop extrusion anchors; blue and yellow highlight conserved and cell-type-specific stripe anchors, respectively. **b**, UMAP projection of ChIP-seq signals showing the co-occupancy by JMJD6 (newly validated binding), H3K27ac, H3K4me3, and RNA Pol II binding profiles. Note that JMJD6 also binds at TAD boundary regions, as presented in Fig. 1c. **c-d**, UMAP visualization of the distance to the nearest transcription start site (TSS, **c**) and stripe scores (**d**) for each CRE in HEK293 cells. **e**, Heatmaps of stripe scores (1 kb resolution, calculated from normalized Micro-C matrix) centered across the five CRE categories. JMJD6–H3K27ac group show the strongest stripe scores, followed by the C–R–Z–J co-occupying group. **f**, High-resolution (1Kb) aggregate peak analysis (APA) visualization of chromatin loops centered on peak pairs across the five CRE categories. **g**, Micro-C contact maps at the *AC005307.3* locus in HEK293T cells showing loss of cohesin loops upon *JMJD6* knockout. **h-i**, Knockout of JMJD6 leads to decreased strength of stripe signals in JMJD6-occupied regions (JMJD6–H3K27ac group and C–R–Z–J co-occupying group). **j-m**, Scatter plot and APA visualization comparing loop strength between wild-type and JMJD6-KO cells at JMJD6–CTCF co-occupied loop sites (**j-k**) and JMJD6-anchored E–P/P–P loop sites (JMJD6–H3K27ac, **l-m**).

To quantify stripe strength genome-wide at high resolution, we developed a stripe score by aggregating Micro-C interaction signals along the diagonal, and followed by localized normalization (Methods). We found stripes are consistently strong at JMJD6-bound sites, with the strongest scores at the enhancer-promoter regions (JMJD6–H3K27ac co-occupied sites), followed by the C–R–Z–J co-occupied boundaries (Fig. 4d-e; Fig. S9c-g). Other sites, although occupied by CTCF, ZNF654, and cohesin, showed no or minimal stripe signals (Fig. 4c-e; Fig. S9c-g).

To further distinguish JMJD6’s transcription-linked functions from classical extrusion anchors, we quantified loop strength across all pairwise combinations of five peak groups (Fig 4f and S9h-j). As expected, loops anchored at C–R–Z–J sites showed the strongest loop-extrusion traces directional to intra-TAD interactions, whereas JMJD6–H3K27ac sites exhibited focal contacts flanked by bidirectional stripes consistent with E–P and P–P loops (Fig. 4f and S9h-j). Together, these results suggest that JMJD6 may participate in both cohesin-mediated extrusion conformation and transcription-associated chromatin interaction networks.

### JMJD6 strengthens stripe and loop interactions

To determine how JMJD6 contributes to mammalian 3D genome organization, we generated JMJD6-KO cells in HEK293 and HCT116 (Fig. S10a-b). Subsequent Micro-C evaluation revealed that, unlike ZNF654-KO that weakens TAD boundary strength and thus increasing inter-TAD interactions, JMJD6-KO cells displayed selective weakening of stripes and focal loops (Fig. 4g and S10). We observed strong reduction in stripe signals at JMJD6–H3K27ac sites and moderate reduction at C–R–Z–J anchors, whereas regions without JMJD6 binding remained unchanged (Fig. 4h-i).

Genome-wide loop analysis revealed that JMJD6 contributes to maintaining the strength of both extrusion-linked and transcription-linked contacts. Upon JMJD6 knockout, the cohesin-extrusion-associated loops and E–P/P–P-loops (JMJD6–H3K27ac) were globally weakened, with 2,337 (22%) and 1,512 (28%) of each category respectively, whereas only ∼1-2% loops were strengthened (Fig. 4j-m). These patterns were consistently observed upon perturbing JMJD6 in HCT116 cells (Fig. S10e-h). Together, these results support a dual role for JMJD6 in reinforcing stripe structures and loop strength at both transcription-associated contacts and extrusion-linked TAD boundaries.

### JMJD6 interacts with BRD2 and BRD4 to strengthen stripe and loop anchors

To investigate potential mechanisms, we performed quantitative ChIP-seq to compare the chromatin factor stability upon JMJD6 knockout. We found that weakened loops frequently coincided with the lost or weakened binding of RAD21 or CTCF at one loop anchor, suggesting that JMJD6 supports cohesin occupancy at a subset of sites asymmetrically (Fig. 4g and S11a-c). Inversely, JMJD6 overexpression promoted RAD21 binding at JMJD6-bound regions (Fig. S11d-e). However, JMJD6 does not directly interact with CTCF or cohesin (Fig. S11f). Previous results suggest that JMJD6 interacts with BRD4 to act as an anti-pause factor for RNA Pol II^36^. Co-immunoprecipitation experiments confirmed the interaction between JMJD6 and BRD4(*34*, *35*), and surprisingly also with BRD2 (Fig. S11g). Previous results indicate that BRD2 tends to bind at TAD boundaries, while BRD4 binds more at enhancer/promoter regions^36,44–46^. Their competitive recruitment of JMJD6 may explain the dual function of strengthening the stripe/loop anchors at TAD boundaries and enhancer/promoter regions, respectively.

Given JMJD6’s strong enrichment at promoter-associated stripes, we examined its impact on transcription. RNA-seq analyses upon JMJD6 knockout revealed widespread downregulation of gene expression, which concurrently also showed pronounced loss of chromatin stripe signals at the promoters (Fig. S11h-i). For example, at *ZNF726* gene whose promoter was bound by JMJD6 to establish a transcription-associated stripe, JMJD6 knockout leads to the depletion of such promoter stripe, and consequently a near-complete loss of gene expression (Fig. S11j-l). These results demonstrate that JMJD6 contributes to transcriptional regulation by stabilizing promoter-centered 3D structures.

## DISCUSSION

In this study, we introduce an *in silico* screen approach that uses a multimodal deep learning model to predict how local genomic perturbations reshape 3D genome architecture. This approach enables genome-wide characterization of structural elements at a resolution and scale not achievable experimentally. By integrating *in silico* screen predictions with chromatin-factor maps, we uncovered two previously uncharacterized proteins – ZNF654 and JMJD6 – that play key functions in shaping mammalian 3D genome architecture. These discoveries refine longstanding models of TAD boundary specification and fine-scale chromatin interactions, while also raising new mechanistic questions.

Our results suggest an updated model of TAD boundary and fine-scale chromatin interaction specification. In this model, CTCF binds at its widespread recognizable genomic sites to provide positional potential, but ZNF654 work together with CTCF at a subset of sites to demarcate the genome into TADs. ZNF654 directly interacts with CTCF, likely acting as an architectural protein complex to reinforce CTCF-anchored TAD boundary insulation and impede cohesin-mediated loop extrusion. This updated model explains why CTCF motif strength, motif clustering, or chromatin accessibility fail to independently predict boundary formation: CTCF binding is necessary but not sufficient, and ZNF654 provides an additional architectural determinant. Further biochemical and structural biology analyses will be required to clarify how ZNF654 work together with CTCF and cohesin during loop extrusion.

In contrast to ZNF654’s TAD boundary-specific function, JMJD6 performs dual architectural and regulatory functions that, to our knowledge, represents the first factor reported to bridge two fundamental layers of genome organization: TAD boundary architecture and transcription-associated chromatin stripes at enhancer–promoter sites. JMJD6 supports cohesin occupancy at a subset of boundary and regulatory anchors, likely through interactions with BRD2 and BRD4, which preferentially bind at TAD boundaries and enhancer–promoter regions, respectively^36,44–46^. JMJD6 has a debating enzymatic function as histone arginine (H3R2 and H4R3) demethylase and lysine hydroxylase^33–35,37^. Whether JMJD6’s enzymatic function contributes to its anchor-defining function – and what exactly is the substrate – remains unknown and requires future studies.

Finally, the discovery of ZNF654 and JMJD6 as novel regulators of chromatin architecture offers insight into the evolutionary origins of 3D genome organization. Although CTCF and cohesin are conserved across metazoans, architectural regulation through cohesin-mediated loop extrusion becomes a dominant mode of genome folding in vertebrates^47–50^. Interestingly, ZNF654 is a vertebrate-specific C2H2 zinc-finger protein, suggesting that its emergence may have enabled or reinforced the establishment of robust loop extrusion and TAD boundary architecture during vertebrate evolution. In contrast, JMJD6 is deeply conserved across metazoans, with human and *Drosophila* JMJD6 proteins sharing >70% sequence identity, far exceeding that of CTCF (42%) or cohesin subunits (∼43-54%). This deep conservation suggests that transcription-coupled focal interactions predate the evolution of vertebrate TAD boundary formation through loop extrusion^48^. These findings raise the possibility that cohesin-mediated loop extrusion mechanism were later integrated with the ancient transcription-associated chromatin interactions to accommodate the expanding genome sizes and regulatory complexity.

## METHODS

### Experimental Methods

#### Cell culture

HCT116 (ATCC, CCL-247) were cultured in McCoy’s 5A medium (ThermoFisher, 16600108) supplemented with 10% FBS (ThermoFisher, 16140071), 1x Penicillin-Streptomycin-Glutamine (100X, ThermoFisher, 10378016). HEK293T cells (ATCC, CRL-3216) were cultured in medium consisting of high glucose DMEM (ThermoFisher, 11995073) supplemented with 10% FBS (ThermoFisher, 16140071) and 1x Penicillin-Streptomycin (100X, ThermoFisher, 15140163). All cells were maintained at 37 °C in a humidified incubator containing 5% CO2. The cells were passaged every 2 days by dissociation with Trypsin-EDTA (0.25%, ThermoFisher, 25200114). Drosophila Schneider 2 (S2) cells (ThermoFisher, R69007) were cultured in Schneider’s Drosophila Medium (ThermoFisher, 21720001) supplemented with 10% heat-inactivated fetal bovine serum (ThermoFisher, 16140071) and 1% Penicillin-Streptomycin (ThermoFisher, 15140163). Cells were maintained at 30 °C in a non-CO incubator as semi-adherent cultures and passaged every 2–3 days at a density of 2–4 × 10 cells/mL. Prior to use as spike-in material, S2 cells were crosslinked with 1% formaldehyde for 10 minutes at room temperature, quenched with 125 mM glycine, washed in PBS, pelleted, and snap-frozen in liquid nitrogen.

### Construction of mCherry Control Plasmid

To generate a negative-control plasmid lacking the hJMJD6 coding sequence, the EF1A–hJMJD6–T2A–PuroR–CMV–mCherry plasmid (VectorBuilder, VB900083) was used as the starting construct. Three primer pairs were designed to amplify fragments excluding the hJMJD6 region, with overhangs compatible for Gibson assembly. PCR amplification was performed using Q5 High-Fidelity DNA Polymerase with GC enhancer (NEB, M0491) in a 50 μL reaction, with an extension time of 30s/kb. Following PCR, 1 μL of DpnI (NEB, R0176S) was added and the reaction was incubated at 37 °C for 1 h to digest template plasmid, followed by heat inactivation at 80 °C for 20 min. PCR products were purified using a PCR purification kit (QIAGEN, 28104). Fragments were assembled using Gibson Assembly Master Mix (NEB, E2611S) in a 20 μL reaction at 50 °C for 15 min, with each fragment added at 0.02–0.5 pmol depending on length and concentration. A positive control assembly reaction was included. Assembled plasmids were transformed into *E. coli* DH5α, and individual colonies were selected for plasmid extraction (ZymoPURE Plasmid Miniprep Kit, D4019). Final constructs were confirmed by full-length plasmid sequencing (PlasmidSaurus).

#### CRISPR/Cas9-mediated genome editing

All guide RNAs of the CRISPR experiments were designed using the CRISPOR algorithm integrated in the UCSC Genome Browser^51^. Guide RNAs were cloned into the pX459V2.0-HypaCas9 plasmid (AddGene, plasmid 62988). CRISPR/Cas9-mediated genome editing was performed largely according to published protocols^52^. Briefly, the sgRNAs were transfected into cells using TransfeX Transfection Reagent (ATCC, ACS-4005) according to the manufacturer’s protocol. In the case of CRISPR knockin cell lines, donor DNA and pX459-sgRNA were transfected in a 1:1 μg ratio. 24 hours later, cells were selected with 2 μg/mL puromycin (ThermoFisher, A1113803) for another 48 hours. Then, cells were cultured in puromycin-free medium for 1 week, after which individual clones were picked and transferred to a 96-well plate and expanded. The homozygous edited clones were identified by a PCR screen from genomic DNA and confirmed by Sanger sequencing. Genomic DNA was prepared with *Quick*-DNA Miniprep Plus Kit (ZYMO RESEARCH, D4069) according to the manufacturer’s protocol. The sequencing chromatograms were aligned in SnapGene. All sgRNAs, insertion template DNA sequences, and genotyping primers are shown in Supplementary Table 1.

### Cell line generation

*JMJD6* KO cells: WT HEK293T and HCT116 were transfected with sgRNA in pX459 targeting the *JMJD6* exon 1 locus. Knock-out of *JMJD6* was confirmed by genomic DNA PCR, Sanger sequencing and western blot.

*ZNF654* KO cells: WT HEK293T and HCT116 were transfected with two sgRNAs in pX459 targeting the *ZNF654* exon 1 locus. Knock-out of *ZNF654* was confirmed by genomic DNA PCR, Sanger sequencing and RT-PCR.

JMJD6-3XFLAG cells: To generate endogenous JMJD6 C-terminal 3XFLAG-tagged cell line, WT HEK293T and HCT116 were transfected with sgRNA in pX459 targeting JMJD6 stop codon and donor DNA flanked on each side by 700 bp of homology to the *JMJD6* stop codon region. The cell line was confirmed by genomic DNA PCR, Sanger sequencing, and western blot. For genomic DNA PCR, 2 primers external to the homology arm were used.

3XFLAG-ZNF654 cells: To generate endogenous ZNF654 N-terminal 3XFLAG-tagged cell line, WT HCT116 were transfected with sgRNA in pX459 targeting ZNF654 start codon, and donor DNA flanked on each side by 700 bp of homology to the *ZNF654* start codon locus. The cell line was confirmed by genomic DNA PCR, Sanger sequencing, and western blot. For genomic DNA PCR, 2 primers external to the homology arm were used.

HA-ZNF654 cells: To generate endogenous ZNF654 N-terminal HA-tagged cell line, WT HCT116 were transfected with sgRNA in pX459 targeting ZNF654 start codon, and donor DNA flanked on each side by 700 bp of homology to the *ZNF654* locus. The cell line was confirmed by genomic DNA PCR, Sanger sequencing, and western blot. For genomic DNA PCR, 2 primers external to the homology arm were used.

### JMJD6 overexpression

HEK293T cells were seeded into 6-well plates and allowed to adhere overnight. The following day, cells in each well were transfected with either EF1A-hJMJD6-T2A-PuroR-CMV-mCherry or EF1A-T2A-PuroR-CMV-mCherry control plasmids using TransfeX Transfection Reagent (ATCC, ACS-4005) according to the manufacturer’s protocol. After 24 hours, the medium was replaced with fresh medium containing 2 µg/mL puromycin for selection. After an additional 24 hours of selection, cells were dissociated with Trypsin-EDTA (0.05%), and cells from three replicate wells of the same condition were pooled and transferred to a 10 cm^2^ dish containing puromycin-containing medium. The following day, the medium was replaced with standard growth medium, and cells were cultured until reaching ∼80% confluency. Overexpression was confirmed by detecting mCherry fluorescence under a fluorescence microscope and western blotting.

### Co-Immunoprecipitation

Cells were washed once with ice-cold PBS, and 10 × 10^6^ cells were lysed in 1 mL Pierce IP Lysis Buffer (ThermoFisher, 87788) supplemented with 1x Protease Inhibitor Cocktail. Lysates were incubated on ice for 30 minutes with periodic mixing, then centrifuged at 4 °C, 16,000 × g for 10 minutes. Supernatants were collected for protein concentration determination and further analysis. For immunoprecipitation, 5 μg primary antibody was added to protein lysate and incubated at 4 °C with rotation for 2 hours. The primary antibody used for IP included: FLAG (Sigma-Aldrich, F1804), CTCF (Diagenode, C15410210), RAD21 (abcam, ab992), SMC1 (ThermoFisher, A300-055A), SMC3 (abcam, ab9263), STAG1 (abcam, ab4457), STAG2 (abcam, ab4464), BRD2 (Cell Signaling Technology Inc, # 5848), BRD4 (GeneTex, Inc, GTX130586), Goat IgG Isotype Control (ThermoFisher, 31245), Mouse IgG Isotype Control (ThermoFisher, 10400C), Rabbit IgG Isotype Control (ThermoFisher, 02-6102). 25 μL of Dynabeads Protein G Beads (ThermoFisher, 10004D) (for antibodies from mouse and goat) or Dynabeads Protein A Beads (ThermoFisher, 10002D) (for antibodies from rabbit) were resuspended by vortexing for 30 seconds and added to the antibody-protein complex. Samples were incubated at 4 °C with rotation for 12-16 hours. The following day, the tube was placed on a magnetic separator, and the supernatant was removed. Beads were washed three times with cold IP wash buffer (add NaCl to Pierce IP Lysis Buffer, final NaCl concentration: 200 mM) by gentle pipetting, with magnetic separation between each wash. For elution, beads were resuspended in 26 μL Pierce IP Lysis Buffer, 10 μL NuPAGE LDS Sample Buffer (4X, ThermoFisher, NP0007) and 4 μL NuPAGE Sample Reducing Agent (10X, ThermoFisher, NP0004). Heat the sample at 95 °C for 10 minutes. Beads were separated magnetically, and the supernatant containing the eluted protein complex was collected for blotting.

### Western blotting

Cells were washed three times in PBS (ThermoFisher, 10010049), a total of 1 × 10^6^ cells were lysed in 100 µl RIPA Lysis and Extraction Buffer (ThermoFisher, 89900) with 1x Protease Inhibitor Cocktail (100X, ThermoFisher, 87786), incubated on ice for 30 minutes, and centrifuged at 4 °C, 12000 x g for 10 minutes. Supernatant was mixed with 1x NuPAGE LDS Sample Buffer (4X, ThermoFisher, NP0007) and 1x NuPAGE Sample Reducing Agent (10X, ThermoFisher, NP0004). Heat the sample at 95 °C for 10 minutes. Samples were run on NuPAGE Bis-Tris 4–12% Mini Protein Gels (ThermoFisher, NP0322BOX) and transferred on to nitrocellulose membrane (ThermoFisher, IB23002) using the iBlot 2 Gel Transfer Device (ThermoFisher, IB21001). Membranes were blocked in 5% non-fat milk (BIO-RAD, 1706404XTU) in Tris-buffered saline with Tween (TBST) at room temperature for 1 hour. The primary antibody was incubated in TBST at 4 °C for 12 hours. Membrane was washed 3 times with 10 minutes each time, then incubated with secondary antibody in 5% BSA in TBST for 1 hour at room temperature, washed 3 times with 10 minutes each time. Chemiluminescent was detected with SuperSigna West Pico PLUS Chemiluminescent Substrate (ThermoFisher, 34580), recorded on a ChemiDoc MP imaging system (BIO-RAD). Primary antibodies used included anti-JMJD6 (1:500, Santa Cruz, sc-48405), anti-FLAG (1:5,000, Sigma-Aldrich, F1804), anti-CTCF (1:1,000, Diagenode, C15410210), anti-RAD21 (1:1,000, abcam, ab992), anti-SMC1 (1:10,000, ThermoFisher, A300-055A), anti-SMC3 (1:25,000, abcam, ab9263), anti-STAG1 (1:10,000, abcam, ab4457), anti-STAG2 (1:10,000, abcam, ab4464), anti-beta-ACTIN [HRP] (1:5,000, NovusBiologicals, NB600-532H). Secondary antibodies used included Goat anti-Rabbit IgG (H+L) Secondary Antibody, HRP (1:5,000, ThermoFisher, 31460), Goat anti-Mouse IgG (H+L) Secondary Antibody, HRP (1:5,000, ThermoFisher, 31430), Rabbit anti-Goat IgG HRP-conjugated Antibody (1:1,000, bio-techne, HAF017).

### ChIP-seq

A total of 5 × 10^6^ cells were collected in 4 mL PBS, 16% methanol-free formaldehyde (ThermoFisher, 28906) was added to the cell suspension to achieve a final concentration of 1% formaldehyde. Cell suspension was incubated at room temperature for 15 minutes on a nutating mixer. The cross-linking reaction was quenched by adding fresh 2.5 M glycine (Alfa Aesar, 36435) to the cell suspension to achieve a final concentration of 125 mM glycine, and incubated for 10 minutes at room temperature on a nutating mixer. For JMJD6 ChIP–seq, cells were crosslinked using a two-step protocol. Cells were first incubated with 2 mM DSG (ThermoFisher, 20593) for 45 min at room temperature. Without washing, 16% methanol-free formaldehyde was added to a final concentration of 1%, and cells were incubated for 15 min on a nutating mixer. The reaction was quenched with 125 mM glycine for 10 min at room temperature. Cells were washed once with ice-cold PBS, collected, and processed alongside samples crosslinked with the single-step formaldehyde protocol used for other factors. Cells were pelleted by centrifugation at 3,000 rpm for 5 minutes at 4°C, the supernatant was carefully removed in a fume hood. The cell pellet was washed twice with 4 mL 4°C PBS with 1x cOmplete EDTA-free Protease Inhibitor Cocktail (Sigma-Aldrich, 04693159001). After each wash, the cells were spun down at 3,000 rpm for 5 minutes at 4°C. Washed cell pellet was resuspended in 1 mL 4°C PBS and transferred into 1.5 mL DNA low-bind tubes (Eppendorf, 022431021), and centrifuged at 4°C for 3.5 minutes at 4,000 rpm. The supernatant was removed as much as possible. Cell pellets were snap-frozen in liquid nitrogen and stored at-80°C until further use. Frozen crosslinked cell pellets were suspended in cell lysis buffer (CLB, 20 mM Tris pH 8.0, 85 mM KCl, 0.5% NP40) with protease inhibitors, incubated on ice for 10 minutes, then centrifuged at 1,000 x g for 5 minutes. Cell pellets were resuspended for a second time in cell lysis buffer with protease inhibitors, incubated on ice for 5 minutes and centrifuged for 5 minutes at 1,000 x g. The pellets were resuspended in nuclear lysis buffer (NLB, 10 mM Tris-HCl pH7.5, 1% NP40, 0.5% sodium deoxycholate, 0.1% SDS) with protease inhibitors for 10 minutes and subsequently sheared in a sonicator (Branson).

Antibodies used in this study included JMJD6 (Santa Cruz, sc-28348), CTCF (Diagenode, C15410210), RAD21 (Abcam, ab992), FLAG (Sigma-Aldrich, F1804), and HA (Abcam, ab9110). ChIP-seq samples were rotated at 4 °C for 12 hours. Protein A magnetic beads (ThermoFisher, 10002D) were used for antibodies raised in rabbits, protein G magnetic beads ((ThermoFisher, 10004D) were used for antibodies raised in mice. For each reaction, 50 µL of Protein A/G beads were aliquoted into a low-bind tube. Beads were washed twice with three volumes of blocking buffer (0.5% TWEEN, 0.5% BSA) containing protease inhibitors. Each wash was performed by adding blocking buffer, mixing by inverting the tube 180° twice, placing the tube on the magnet, and removing the supernatant. After the final wash, beads were resuspended in 50 µL of ChIP Dilution Buffer (16.7mM Tris-HCl pH8.1, 1.1% Triton X-100, and 167mM NaCl, 1.2mM EDTA, 0.01% SDS) with protease inhibitors per reaction. ChIP samples were removed from rotation, pulse-spun, and 50 µL of pre-blocked beads were added to each sample. Samples were incubated at 4°C for 1 hour with end-over-end rotation. The ChIP-seq samples were removed from rotation, spun briefly and placed on a magnet to isolate the beads. The beads were washed with a series of buffers: low salt RIPA buffer (0.1% SDS, 1% Triton X-100, 1mM EDTA, 20mM Tris-HCl pH 8.1, 140mM NaCl, 0.1% DOC), high salt RIPA buffer (0.1% SDS, 1% Triton X-100, 1mM EDTA, 20mM Tris-HCl pH 8.1, 500mM NaCl, 0.1% DOC), LiCl buffer (250 mM LiCl, 0.5% NP40, 0.5% sodium deoxycholate, 1 mM EDTA, 10 mM Tris-HCl pH 8.1) and finally Low TE buffer (10mM Tris pH 7.5). The Protein A/G beads were then suspended in 50 µl direct ChIP elution buffer (10 mM Tris-Cl pH 8.0, 5 mM EDTA, 300 mM NaCl, 0.1% SDS and 5 mM DTT directly before use) and 8 µl of reverse crosslinking mix (250 mM Tris-HCl pH 6.5, 1.25 M NaCl, 62.5 mM EDTA, 5 mg/ml Proteinase K [ThermoFisher, EO0491], and 62.5 µg/ml RNAse A). The suspended beads were incubated at 65 °C for 3 hours. After incubation, the supernatants were transferred to a clean tube. The DNA was purified by Beckman Coulter AMPure XP beads (Beckman Coulter A63881), eluted, and quantified by Qubit dsDNA HS Assay Kit (Invitrogen Q33231). Illumina library preparation was performed using the NEBNext Ultra II kit (New England BioLabs, E7645).

### Quantitative ChIP-seq (qChIP-seq)

For quantitative ChIP-seq, WT and KO cells were cultured in parallel under identical conditions. For each replicate, 6 million cells were collected and crosslinked with 1% formaldehyde for 10 minutes at room temperature, followed by quenching with 125 mM glycine. During the first wash with CLB buffer, crosslinked Drosophila S2 cells were added at 5% of the human cell number as spike-in material. After chromatin shearing, samples were split into two aliquots and incubated with either CTCF or RAD21 antibody. In both WT and KO samples, 1 ug spike-in antibody recognizing the Drosophila-specific histone variant H2Av (Active Motif, 61686) was added. Immunoprecipitation was performed overnight at 4 °C with rotation. The next day, Protein A and Protein G magnetic beads (mixed at 1:1 ratio) were added to enrich the antibody–chromatin complexes. All remaining steps were performed identically to standard ChIP-seq.

### Micro-C

Micro-C protocol was adapted from the original protocol with minor changes that optimizes sample quality^6,7,53^. sample cells were double-crosslinked to fix protein-protein and protein-DNA interactions using 3mM DSG (disuccinimidyl glutarate, 7.7 A) (ThermoFisher, 20593) for 35 min and 1% formaldehyde (ThermoFisher, 28906) for 10min at room temperature. The reaction was then quenched with 0.375 M Tris buffer pH 7.5 (ThermoFisher, 15567027). Crosslinked cells were washed with 1× cold PBS at a concentration of 1 × 10^6^ cells per ml, centrifuged for 5 minutes at 850 xg at 4 °C. Cell pellet was resuspended in cold PBS and aliquoted into multiple tubes containing 5 × 10 cells per tube. Centrifuge the tubes for 5 min at 850 × g at 4 °C and remove the supernatant. The resulting cell pellets were snap-frozen in liquid nitrogen and stored at −80 °C until use.

### MNase titration

Prepare fresh MB1 buffer (50 mM NaCl, 10 mM Tris, 5 mM MgCl_2_, 1 mM CaCl_2_, 0.2% NP-40, 1xProtease inhibitor cocktail). Cell pellets were thawed on ice and resuspended in complete MB1 buffer at a concentration of 1 × 10^6^ cells per 100 µL in low-retention 1.5-mL tubes. Cells were incubated on ice for 20 minutes, followed by centrifugation at 1,750 × g for 5 minutes at 4 °C. Supernatants were carefully aspirated. Nuclei pellets were washed by resuspension in MB1 buffer at a concentration of 1 × 10^6^ nuclei per 100 µL. Pellets were gently pipetted up and down until fully resuspended, then centrifuged at 1,750 × g for 5 minutes at 4 °C. Supernatants were carefully removed. 20 U/μL Micrococcal nuclease (ThermoFisher, EN0181) was diluted 1:20 with 10 mM Tris-HCl, pH 7.5 (ThermoFisher, 15567027). Nuclei pellets were resuspended in MB1 buffer at a concentration of 1 × 10^6^ nuclei per 100 µL by gentle pipetting. Increasing volumes of the diluted MNase (4, 7, 10, 15, or 20 µL) were added to individual aliquots to perform a titration test, corresponding to increasing enzyme concentrations. Samples were briefly vortexed to mix and incubated at 37 °C for 20 minutes with shaking at 1,000 rpm. Digestion was immediately halted by adding 500 mM EGTA to a final concentration of 4 mM. Samples were incubated at 65 °C for 10 minutes, and then centrifuged at 2,000 × g for 5 minutes at 4 °C. Pellets were resuspended in 150 µL of Reverse Crosslinking Solution (1xTE, 1 mg/ml Proteinase K, 1% SDS, 100 µg/mL RNaseA, 0.2 M NaCl). Cross-links were reversed by incubating the samples at 65 °C for 2 h. Then DNA was purified using the Zymo DNA Clean & Concentrator kit (Zymo Research, D4033). Purified DNA was resolved by agarose gel electrophoresis. The sample exhibiting the optimal digestion pattern - approximately 80–90% mononucleosomes and 10–20% dinucleosomes - was selected for downstream applications.

### MNase digestion

To extract intact nuclei, crosslinked cell pellets (5 × 10^6^ cells per sample) were resuspended in MB1 buffer at a density of 1 × 10^6^ cells per 100 µL. Samples were incubated on ice for 20 minutes to solubilize cell membranes. Nuclei were pelleted by centrifugation and washed once with ice-cold MB1 buffer. Following the wash, nuclei were resuspended in 500 µL MB1 buffer. MNase (pre-determined by titration) was added at the appropriate volume from a 20 U/µL stock. Samples were incubated at 37 °C for 20 minutes in a thermomixer with shaking at 1,000 rpm. The reaction was quenched by the addition of EGTA to a final concentration of 4 mM (bioWORLD, 40520008), followed by heat inactivation at 65 °C for 10 minutes. Digested nuclei were subsequently washed twice with ice-cold MB2 (50 mM NaCl, 10 mM Tris-HCl pH 7.5, 10 mM MgCl_2_, and 100 µg/mL BSA [Sigma-Aldrich, B8667]).

### End repair and labeling

To generate blunt-ended, biotin-labeled DNA fragments suitable for proximity ligation, a series of enzymatic steps were performed on digested nuclei. End-repair reactions were first carried out to phosphorylate 5’ termini and remove 3’ phosphate groups. Samples were incubated at 37 °C for 15 minutes in a reaction containing 50 U T4 Polynucleotide Kinase (New England BioLabs, M0201), 50 mM NaCl, 10 mM Tris-HCl pH 7.5, 10 mM MgCl2, 100 µg/mL BSA, 2 mM ATP (ThermoFisher, R1441), and 5 mM DTT (Sigma-Aldrich, 10197777001) in nuclease-free water, shaking at 1,000 rpm. To generate 5′ overhangs for subsequent end-blunting and biotin labeling, 50 U DNA Polymerase I Klenow Fragment (New England BioLabs, M0210) was added to the same reaction. The mixture was incubated at 37 °C for an additional 15 minutes with shaking at 1,000 rpm. Next, a nucleotide mix was added for end-labeling, consisting of 66 µM each of dTTP (Jena Bioscience, NU-1004), dGTP (Jena Bioscience, NU-1003), biotin-dATP (Jena Bioscience, NU-835-BIO14), and biotin-dCTP (Jena Bioscience, NU-809-BIOX) in 1× T4 DNA Ligase Buffer supplemented with 100 µg/mL BSA. This reaction was incubated at room temperature for 45 minutes with intermittent mixing (1 minute of shaking, followed by 3 minutes still, repeated). Reaction was quenched by adding EDTA to a final concentration of 30 mM (Invitrogen, 15575020), followed by heat inactivation at 65 °C for 20 minutes. The biotin-labeled, end-repaired nuclei were washed once with ice-cold MB3 buffer (50 mM Tris-HCl pH 7.5, 10 mM MgCl2, and 100 µg/mL BSA).

### Proximity ligation and removal of unligated biotin

Proximity ligation was carried out by incubating biotin-labeled chromatin in a 500 µL ligation reaction containing 10,000 U T4 DNA Ligase (New England BioLabs, M0202), 1× T4 DNA Ligase Buffer, and 100 µg/mL BSA, in nuclease-free water. Samples were incubated at room temperature for 2.5 h with gentle mixing. To remove biotinylated nucleotides from unligated DNA ends, samples were treated with 1,000 U Exonuclease III (New England BioLabs, M0206) in 1× NEBuffer #1. Reactions were incubated at 37 °C for 15 minutes with intermittent mixing (1 minute of shaking, followed by 3 minutes still, repeated).

### DNA purification, size-selection and library construction

Samples were supplemented with 1% SDS (Sigma-Aldrich, L3771), 2 mg/mL Proteinase K, 250 mM NaCl, and 100 µg/mL RNase A (ThermoFisher,EN0531), then incubated at 65 °C overnight (12-16 h). DNA was purified using the Zymo DNA Clean & Concentrator kit according to the manufacturer’s instructions. Dinucleosome-sized DNA fragments (250–400 bp) were isolated via electrophoresis on a 1.5% agarose gel (VWR, 97062), excised, and purified using the Zymo Gel Purification Kit (Zymo Research, D4008). DNA concentrations were measured with the Qubit 1× dsDNA High Sensitivity Assay (Invitrogen, Q33231). Biotinylated, ligated DNA fragments were then isolated using Dynabeads MyOne Streptavidin T1 (Invitrogen, 65601). 1 mL of 1× Tween Binding and Wash (TBW) buffer (1 M NaCl, 5 mM Tris-HCl pH 7.5, 500 µM EDTA, 0.1% Tween-20 [Sigma-Aldrich, P8074]) was added to 25 µL T1 beads, gently mixed on a nutator for 2 minutes, followed by magnetic separation for 60 s. The supernatant was carefully removed. Then, beads were resuspended in 150 µL of 2× Binding and Wash buffer (2 M NaCl, 10 mM Tris-HCl pH 7.5, 1 mM EDTA). In parallel, the volume of purified DNA was brought to 150 µL. Pre-washed beads were then added to each sample, and incubated at room temperature with gentle rotation for 20 minutes. Samples were washed twice with 1× TBW Buffer. For each washing, samples were briefly centrifuged in a benchtop microcentrifuge, and 950 µL of 1× TBW buffer was added. Tubes were inverted and shaked at 1,200 rpm for 5 minutes at room temperature. Following brief centrifugation, samples were placed on a magnetic stand for 60 s, and the supernatant was carefully removed. Bead-bound DNA was subsequently used for library preparation with the NEBNext® Ultra™ II DNA Library Prep Kit for Illumina® (New England BioLabs, E7645), following the manufacturer’s protocol. All steps were performed as directed by the manual, with the exception that end-repair and A-tailing, adapter ligation incubations were performed with intermittent shaking (1 minute shaking followed by 3 minutes still, repeated). Bead-bound samples were washed to remove residual enzymes and unligated adapters. Samples were incubated with 950 µL of 1× TBW buffer, inverted, and shaked at 1,200 rpm for 3 minutes at room temperature. Tubes were briefly centrifuged, placed on a magnetic stand, and the supernatant was carefully removed. Repeated wash step once. Beads were resuspended with 20 µL elution buffer for downstream applications. To determine the minimum number of PCR cycles required for adequate input material for sequencing, a test amplification (11 and 14 cycles) was performed using 1 µL sample to quantify yield. The test PCR products were analyzed by agarose gel electrophoresis, and the yield was quantified using ImageJ. All library amplifications incorporated sequencing indices from the NEB Multiplex Oligos for Illumina Primer Set 1–4 (New England BioLabs, E7335). Following amplification, the original T1 bead-bound samples were removed. The amplified libraries were then purified using 0.8 x AmPure XP beads (Beckman Coulter, A63880) to remove adaptor dimers, primers, and other contaminants. Purified libraries were quantified using Bioanalyzer and Qubit to determine the size distribution and final concentrations.

### In situ HiC

*In situ* Hi-C was performed on 5 million HEK293 cells using the Arima-HiC Kit (Arima Genomics, A510008) according to the manufacturer’s protocol. Briefly, cells were crosslinked with 1% formaldehyde for 10 minutes at room temperature and quenched with glycine. Nuclei were isolated and chromatin was digested with a restriction enzyme mix provided in the kit. Following end-repair and incorporation of biotinylated dNTPs, proximity ligation was carried out in intact nuclei to capture spatially adjacent DNA fragments. Crosslinks were reversed, and DNA was purified. Final libraries were constructed using the NEBNext Ultra II DNA Library Prep Kit(New England BioLabs, E7645) for Illumina sequencing.

### RNA-seq

Total RNA was extracted using the RNeasy Mini Kit (Qiagen, 74004) according to the manufacturer’s instructions. Two biological replicates were prepared for each condition, with replicates derived from independent clones where applicable. RNA sequencing libraries were constructed using the NEBNext Ultra II Directional RNA Library Prep Kit (NEB, E7765L) following the rRNA depletion workflow.

### Computational analysis

#### *In silico* genetic screen with C.Origami

We performed an *in silico* screen to identify sequence elements that exert strong influence on 3D genome architecture. For each chromosome, we scanned the genome with a 2-Mb window and introduced a 1-kb deletion centered within the window. Both the unperturbed and perturbed inputs were predicted using C.Origami^30^, and the resulting contact maps were compared to compute an impact score, defined as the pixel-wise mean absolute difference between the two predictions. The window was then shifted by 1 kb, generating a continuous 1-kb–resolution impact profile across the genome. To reduce computational load, ISGS was performed on ten representative chromosomes (chr5, 7, 8, 11, 12, 14, 15, 19, 20 and 22). To identify impactful loci, we calculated a peak score for each position based on the local range across a three-bin window and selected the top 1% as high-impact elements. Next, we integrated binding profiles of all chromatin-associated proteins (CAPs) from the ReMap2022^31^ database and ranked each factor by the enrichment of its binding sites within high-impact loci.

### ChIP-seq analysis

Paired-end reads were quality-trimmed using Trim Galore (v0.6.7) with parameters-q 25 --paired --phred33-e 0.1 --clip_R1 10 --clip_R2 10 --length 36 --stringency 3. Sequencing quality was assessed using FastQC (v0.11.9). Trimmed reads were aligned to the human reference genome (hg38) using Bowtie2 (v2.4.5)^54^ with default settings, and aligned reads were converted to BAM format using samtools (v1.15.1)^55^. PCR duplicates were removed using Picard (v2.26.10) with parameters MarkDuplicates REMOVE_DUPLICATES=true, and deduplicated BAM files were indexed. Normalized coverage tracks were generated using bamCoverage (deepTool v3.5.1)^56^ with parameters --binSize 25 --smoothLength 75 --normalizeUsing CPM –centerReads--extendReads, excluding reads mapping to mitochondrial DNA (chrM). All tracks were CPM-normalized while datasets intended for quantitative comparison of ChIP-seq (qChIP-seq) signals were processed without CPM normalization. Peak calling was performed using MACS2 (v2.2.7.1)^57^ with the callpeak function in paired-end mode (-f BAMPE). For JMJD6 and ZNF654 in this study, peaks were called with a q-value cutoff of 0.05. For all other factors, peaks were called using q-value threshold of 0.01. To generate a high-confidence HCT116 HA-ZNF654 peak set for downstream analysis given the high background signal, HA-ZNF654 signal intensity was ranked across HCT116 CTCF peaks in descending order, and the top 30,000 peaks were selected. This set was then unioned with peaks called by MACS2, and overlapping regions were merged into contiguous intervals spanning the minimum start and maximum end coordinates. Peaks from all factors overlapping blacklisted regions were excluded prior to downstream analysis. Blacklisted regions were defined as the union of non-specific IgG ChIP-seq peaks and unmappable regions identified from Micro-C data.

### Quantitative ChIP-seq (qChIP-seq) analysis

Reads were aligned to a concatenated human (hg38) and Drosophila melanogaster (dm6) genome using Bowtie2 (v2.4.4) with parameters --no-mixed --no-discordant. During concatenation, chromosome names were prefixed with “hg38_” or “dm6_” to facilitate species-specific separation after mapping. After alignment, secondary alignments were removed by filtering out reads containing the XS: tag, and unmapped reads were discarded using samtools view-F4 (v1.13). Duplicate reads were removed using sambamba markdup --remove-duplicates (v0.8.0). Mapped reads were then separated by species based on chromosome prefixes, and chromosome names were restored to their original formats.

For each pairwise comparison (*e.g.* A vs B), spike-in normalization was carried out using Drosophila dm6-aligned reads as an internal reference. The sample with the lower dm6 spike-in count was used as the normalization baseline, and human (hg38) reads in the other sample were down-sampled accordingly to ensure consistent spike-in levels across conditions. Specifically, the down-sampled human reads for each sample were calculated as:

Where **dm6*_A_***, **dm6*_B_*** are the number of dm6-aligned reads in sample A and B respectively, and **hg38_sample_** is the number of human-aligned reads in the corresponding sample.

Down-sampling was performed using sambamba view --subsampling-seed=123^58^. For down-sampled BAM files, coverage tracks were generated using bamCoverage (deepTools v3.5.1) with parameters --binSize 25 --smoothLength 75 --centerReads --extendReads, excluding mitochondrial reads (chrM). No additional normalization was applied to preserve the spike-in–adjusted quantitative signal. The resulting bigWig files were used for all downstream analyses.

### ChIP-seq peak overlap and annotation

ChIP-seq peaks for CTCF, RAD21, ZNF654, JMJD6 and H3K27ac were individually filtered to remove blacklisted regions (hg38) using bedtools intersect-v (v2.27.1)^59^. Combinatorial categories of those peak regions, including C-R-Z-J, C-R-Z, C-R, CTCF-only, JMJD6-H3K27ac, were defined through sequential overlaps computed using bedtools intersect-wa combined with exclusion operations (-v). Three-way intersections were visualized with the venn3 function (matplotlib-venn v0.11.7) based on the actual counts of overlapping peaks. Four-way intersections were manually adjusted for proportional representation, as current visualization tools could not accurately preserve the relative sizes of overlapping and non-overlapping regions. Genomic elements including coding exons (CDS), 5’ untranslated regions (5’UTR), 3’ untranslated regions (3’UTR), and introns were downloaded from the UCSC Genome Browser (knownGene V46 track, GRCh38/hg38 assembly). Transcription start sites (TSSs), H3K27ac peaks, and CTCF peaks were also collected and merged with these annotations to generate a comprehensive genomic feature set. Peaks from ChIP-seq datasets were intersected with annotated features using bedtools intersect-wa-wb. To resolve cases where a peak overlapped multiple features, each peak was assigned to a single category based on a predefined priority: CTCF > TSS > Enhancer (H3K27ac) > CDS > 5′UTR > 3′UTR > Intron. Only the annotation with the highest priority was retained per peak. Peaks without overlap were classified as intergenic. Peak distributions across feature categories were summarized and visualized as a stacked bar plot using a custom Python script, preserving the proportions and applying consistent color coding across annotations.

### UMAP embedding of *cis*-regulatory element features

Candidate *cis*-regulatory elements (cCREs) were defined by merging peaks from ChIP-seq and ATAC-seq datasets, excluding regions overlapping the hg38 blacklist regions. The ChIP-seq datasets included profiles for CTCF, RAD21, ZNF654, JMJD6, H3K27ac, H3K4me3, and RNA polymerase II. Overlapping ChIP-seq and ATAC-seq peaks were merged into unified intervals using bedtools (v2.30.0). For each cCRE, mean signal intensities from each ChIP-seq and ATAC-seq dataset were extracted from bigWig files using pyBigWig (v0.3.18). The resulting signal matrix was log-transformed (log1p) and z-score normalized across regions. Principal component analysis (PCA) was performed for dimensionality reduction, followed by UMAP projection using scanpy (v1.9.3)^60^. The resulting UMAP embedding was preserved for downstream overlay of additional genomic and epigenomic features. Specifically, ChIP-seq enrichment scores for additional factors, Micro-C derived metrics (e.g., insulation scores and z-score–normalized stripe strength), and genomic features such as distance to the nearest transcription start site (TSS) were mapped onto the same UMAP coordinates to enable integrated visualization and comparative analysis of cCRE properties. cCREs were clustered using the Leiden algorithm (resolution = 1.0), and enrichment patterns were visualized across the UMAP space.

To evaluate the spatial relationships between cCREs and transcriptional regulation, we quantified the genomic distance from each cCRE center to its nearest TSS. For genes with multiple annotated TSSs, we selected the one exhibiting the strongest local stripe signal (within a ±500 bp window) as the representative TSS, reflecting putative regulatory engagement. To constrain the analysis to biologically plausible long-range interactions and reduce computational load, candidate TSSs were restricted to a ±4 Mb window centered on each cCRE. This range encompasses most loop-and stripe-mediated cis-regulatory interactions, particularly those involving architectural factors such as CTCF and RAD21. The distance to the nearest TSS within this window was calculated for each cCRE and projected onto the UMAP embedding for integrative visualization and comparative analysis of regulatory potential.

### Micro-C and *in situ* Hi-C data analysis

Paired-end reads generated by Illumina sequencing were aligned to the UCSC hg38 reference genome using BWA-MEM (v0.7.17)^61^ with the options-SP5M to enable split-read mapping and preserve pairing information. Aligned reads were parsed using pairtools (v0.3.0) with the following options: pairtools parse --add-columns mapq --drop-sam --drop-seq. Parsed reads were then sorted using pairtools sort, and valid read pairs of type UU, UR and RU were retained using pairtools select. Biological replicates from the same condition were merged at this step, and PCR duplicates were marked and removed using pairtools dedup with the --mark-dups option. To ensure comparability across conditions, wild-type and knockout samples for both ZNF654 and JMJD6 were sequenced to similar depths. To ensure equal sequencing depth across samples, deduplicated contact pairs were down-sampled to match the contact count of the sample with fewer reads. This was done by separating the pairtools header from the deduplicated contact pairs file, randomly shuffling the interaction records, and retaining the desired number of contacts. Down-sampled pairs were concatenated with the original header and reprocessed using pairtools sort. The deduplicated down-sampled contact pairs were converted to the.hic format using Juicer tools(v1.22.01)^62^ with multi-resolution binning at 1000000, 250000, 100000, 50000, 25000, 10000, 5000, 2000, 1000, 500, and 200 bp, followed by matrix balancing using the addNorm function. In parallel, cool files were generated using cooler (v0.9.1)^63^ via the cload pairs function at 50 bp resolution. Multi-resolution contact matrices were then generated using cooler zoomify with bin sizes ranging from 50 bp to 250 kb and balanced using the --balance option.

### Analysis of contact probability decay and slope

Contact probability decay curves, P(s), were generated from 1-kb resolution balanced Hi-C matrices using cooltools (v0.7.1)^64^. To compute genome-wide P(s) profiles, intra-chromosomal contact frequencies were averaged across chromosome arms, defined based on hg38 chromosomal coordinates and centromere positions retrieved from the bioframe package. For each sample, cis-expected contact frequencies were calculated using the expected_cis function with a Gaussian smoothing sigma of 0.1, and aggregate-smoothed contact frequencies were extracted. Distances below 2 kb were masked to minimize the impact of short-range noise. After ensuring one data point per genomic distance, P(s) curves were plotted on log-log axes. To characterize the scaling behavior of contact decay, the local slope of the P(s) curve was computed by taking the first derivative in log-log space using numpy.gradient. Slope profiles were visualized as a function of genomic separation on a semi-logarithmic scale.

### Insulation score

Insulation scores were calculated using cooltools on balanced contact matrices at 5-kb and 1-kb resolutions. For each resolution, a sliding window corresponding to 25 bins (125 kb and 25 kb, respectively) was applied. To minimize contributions from adjacent bins, interactions within three diagonals were ignored during calculation. Resulting insulation profiles were exported as BigWig files for visualization and further used for downstream analysis.

### Quantitative analysis of Micro-C loops

Loops were called at resolutions of 10 kb, 5 kb, 2 kb, and 1 kb using Mustache (v1.0.1)^65^ with the parameters-pt 0.1-st 0.88 --octaves 2. Loop calls from multiple resolutions were merged to create a unified set of high-confidence loops. Loop coordinates from each resolution were first tagged with a unique loop identifier and concatenated, then sorted using bedtools. To remove redundancy, overlapping loops were filtered using a custom Python script. Briefly, loops were ranked by size and, for each overlapping group (where both anchors overlapped), only the highest resolution loop was retained to avoid redundancy from multi-resolution calling. To compare loop strengths between WT and KO samples, Micro-C interaction values were extracted from KR-normalized.hic files using straw (v0.1.0)^62^, with loop regions queried at dynamically selected resolutions (1 kb, 2 kb, 5 kb or 10 kb) based on loop size to ensure appropriate resolution for each interaction. For each loop, the sum of observed-over-expected (OE) interaction values was calculated across the two anchor regions. Log-transformed values were computed to quantify interaction strength, with appropriate handling for zero values.

Loops were classified as JMJD6-bound or ZNF654-bound if either anchor overlapped JMJD6 or ZNF654 binding sites. JMJD6/ZNF654-associated loops were further divided into CTCF-associated or enhancer–promoter loops based on the presence of CTCF or enhancer–promoter elements at either anchor. Loops with a log fold change greater than 0.5 or less than –0.5 between WT and KO samples were defined as significantly differential and visualized in scatter plots. These loops were further validated by aggregate peak analysis (APA) using coolpup (v1.1.0)^66^ on 1 kb resolution contact maps in.mcool format. APA profiles were visualized using plotpup (v1.1.0) with asymmetry preserved.

### Chromatin stripe analysis

Chromatin stripe signals were calculated using both raw and ICE-normalized Micro-C contact matrices at 1 kb, 500 bp, and 200 bp resolutions. All computations were performed on the full symmetric contact matrix. For each bin along the main diagonal, interaction values were summed along an off-diagonal axis deviating from the 45° diagonal of the Hi-C matrix, extending 1 Mb upward and 1 Mb downward from the focal bin. Each flanking region was defined as a one-bin-wide diagonal band extending from the central bin, excluding three pixels along the main diagonal to minimize short-range interaction noise. Stripe signals were computed genome-wide by sliding this window along the diagonal, and the final signal for each bin was defined as the sum of upstream and downstream interactions.

To correct for local variation in interaction density and mitigate noise arising from data sparsity, a local z-score normalization was applied. This normalization also enabled direct comparison of stripe signals between different samples with similar sequencing depth. Specifically, for each bin, a surrounding genomic window around the central bin (500 × resolution) was used to calculate the local mean (μ) and standard deviation (σ) of total interaction signals. The normalized z-score was then computed as:

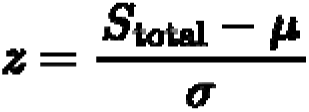

where ***S*_total_** represents the sum of upstream and downstream interactions at each bin. Bins with undefined or zero variance within the local window were assigned a z-score of zero. Normalized stripe profiles were exported as BigWig tracks for genome-wide visualization and downstream analysis.

### Visualization of Genomic Features

To integrate 3D genome contact patterns with regulatory and transcriptional landscapes, we generated unified genomic region plots that display Micro-C contact maps, ChIP-seq and RNA-seq profiles, CTCF motif orientations, and gene annotations. Balanced Micro-C contact matrice were extracted using cooler at selected resolutions for specified genomic regions of interest. Zero and NaN entries were masked and rendered in a soft blue background (RGB: 0.78, 0.84, 0.93) to distinguish missing values from valid interactions. Contact maps for WT and KO samples were visualized using matplotlib (v3.10.3) with the’fall’ colormap, and the maximum color intensity (vmax) was set to the 95th percentile of non-zero contact values, with manual adjustment for certain regions to improve visual contrast. Differential contact maps (KO − WT) were computed and visualized using the’seismic’ diverging colormap, with color limits (vmin and vmax) set at the 98th percentile of absolute differences.

All signal tracks - including ChIP-seq, RNA-seq, insulation scores, and z-score normalized stripe strengths - were smoothed and displayed with consistent binning and unified y-axis scaling, enabling direct visual comparison across chromatin structure, regulatory activity, transcriptional output, and gene organization. CTCF motif orientations were indicated with strand-specific arrows, and gene annotations (in BED12 format) were retrieved from GRCh38.p14 with transcript isoforms collapsed for clarity. All elements were assembled into a unified layout using customized scripts based on the CoolBox API (v0.3.9)^67^.

### Pileup analysis of chromatin interaction maps

Aggregated ChIP-seq signal profiling around genomic features was performed using computeMatrix (deepTools v3.5.1) in reference-point mode, centered on each feature with a ±1 kb flanking window unless otherwise specified. Zeros and missing values were excluded using the --skipZeros and --missingDataAsZero options. Heatmaps were generated using plotHeatmap (deepTools), where each ChIP-seq dataset was visualized with a two-color scale: white indicating background and a distinct color highlighting signal enrichment. Insulation scores and stripe strengths were visualized with diverging color maps to differentiate signal polarity.

Aggregate pileup analyses of Micro-C data were performed using coolpup.py (coolpuppy v1.1.0). Balanced contact matrices were extracted, and contact enrichment around ChIP-seq defined genomic features was calculated using the --local option. Unless otherwise specified, a 250 kb flanking window was applied on each side of the feature center. Aggregated contact maps were visualized using <u>plotpup.py</u> (coolpuppy), with diverging colormaps adjusted to highlight relative enrichment or depletion around the features of interest.

### Loop enrichment analysis

Candidate chromatin interaction pairs were defined between genomic features based on BED intervals using a custom Python script. Genomic regions corresponding to distinct feature categories, such as C-R-Z-J, C-R-Z, C-R, CTCF-only, and JMJD6-H3K27ac, were used as input sets. All possible intra-category and inter-category region pairs were first enumerated. For each pair, the genomic distance between their peak centers was calculated, and only those between 100 kilobases (kb) and 2 megabases (Mb) were retained. The lower bound was set to exclude short-range background contacts, while the upper bound reflects the typical span of cohesin-mediated loops and matches commonly used parameters in HiChIP and related assays^68^. The distance-filtered pairs were saved in BEDPE format and used for downstream aggregate peak analysis (APA) to quantify Micro-C contact enrichment at 5-kb and 1-kb resolutions.

### Quantifying local density of CTCF peaks across genomic windows

Peaks from different groups (C-R-Z-J, C-R-Z, C-R, and CTCF only) were merged into a unified dataset while retaining group annotations. To assess the degree of spatial clustering of CTCF binding on the genome, the number of neighboring peaks within symmetric genomic windows of varying sizes (±1 kb, ±3 kb, ±5 kb, ±20 kb, ±50 kb, and ±100 kb) was quantified for each peak using a custom Python script. Peaks from all groups were included in the neighborhood counts; however, the query peak itself was excluded by subtracting one from the total number of overlaps.

### CTCF Motif analysis

CTCF motifs within peaks from each CTCF group were scanned using gimme scan (gimmemotifs v0.18.0)^69^ with a motif match cutoff of 0.99 and the JASPAR 2024 CTCF position weight matrix (PWM)^70^. The resulting motif predictions were intersected with annotated CTCF motifs from the UCSC Genome Browser. Motif scores were calculated as –log (p-value) × 100 for each overlapping site. The distribution of motif scores across different groups was visualized using violin plots to facilitate comparison of CTCF motif strength and positional conservation.

### RNA-seq analysis

Raw RNA-seq reads were aligned to the human reference genome (GRCh38/hg38) using STAR (v2.7.2a)^71^ with default parameters. Strand-specific read counts were summarized into a count matrix for downstream analysis. Principal component analysis (PCA) was conducted using the top 500 most variable genes across all samples to assess transcriptomic variance. Differential gene expression analysis was performed using DESeq2 (v1.40.2)^72^. Genes with an absolute log (fold change) > 0.5 and an adjusted P-value < 0.05 were considered significantly differentially expressed. The TSSs of significantly upregulated and downregulated genes were extracted and used as anchors for aggregate pile-up analysis on 1 kb resolution Micro-C contact maps, enabling comparative assessment of local chromatin architecture in wild-type (WT) and *JMJD6* knockout (KO) cells. Bar plots of selected gene expression level were generated using normalized counts from DESeq2 (size-factor corrected) for selected genes, and p-values were derived from the Wald test implemented within DESeq2 and annotated directly onto the plots.

### Genomic Datasets

| Protein | Cell line | Data source | Accession code |
| --- | --- | --- | --- |
| ZNF654 | HEK293 | ENCODE | ENCSR504VDV |
| JMJD6 rep1 | HEK293 | Liu et al, 2013<sup>36</sup> | GSM1249879 |
| JMJD6 rep2 | HEK293 | Liu et al, 2013 | GSM1249880 |
| JMJD6 | Hela | Liu et al, 2013 | GSM1249905 |
| JMJD6 | Mouse Th17 | Ciofani et al., 2012<sup>73</sup> | GSM1004840 |
| RNA Pol II | HEK293 | Liu et al, 2013 | GSM1249894 |
| H3K27ac | HEK293 | Liu et al, 2013 | GSM1249889 |
| RNA Pol II | HCT116 | ENCODE | ENCSR000EUU |
| H3K27ac | HCT116 | ENCODE | ENCSR661KMA |
| CTCF | Hela | Pflueger et al., 2018<sup>74</sup> | GSM2872304 |
| RAD21 | Hela | Li et al., 2019<sup>75</sup> | GSM2990425 |
| SMC3 | Hela | Stocsits et al., 2020<sup>76</sup> | GSM4106811 |
| SMC3 | Mouse Th17 | Cheung et al., 2017<sup>45</sup> | GSM1902312 |
| CTCF | Mouse Th17 | Ciofani et al., 2012<sup>73</sup> | GSM1004804 |
| NIPBL | HCT116 | Rao et al., 2017<sup>77</sup> | GSM2809637 |
| SMC1 | HCT116 | Nanavaty et al., 2020<sup>78</sup> | GSM3790846 |
| SMC3 | HEK293 | Parenti et al., 2020<sup>79</sup> | GSM3463658 |
| STAG2 | AML3 | Smith et al., 2020<sup>80</sup> | GSM3032902 |
| STAG1 | K562 | ENCODE | ENCSR153HNT |
| ZNF574 | HEK293 | PMID: 27852650 | GSM2026820 |
| PRKDC | Fibroblast | PMID: 25057768 | GSM1340090 |
| TRIM22 | GM12878 | ENCODE | ENCSR637QAM |
| CXXC4 | HEK293 | Ko et al., 2013<sup>81</sup> | GSM1054009 |
| ZNF660 | HEK293 | ENCODE | ENCSR283DOU |
| ZSCAN30 | HEK293 | ENCODE | ENCSR768VNZ |
| SUPT5H | K562 | ENCODE | ENCSR894CGX |
| PATZ1 | HEK293 | ENCODE | ENCSR966ULI |
| MAZ | HEK293 | ENCODE | ENCSR290SSQ |
| ZNF263 | HEK293 | ENCODE | ENCSR000EVD |
| ZNF341 | HEK293 | ENCODE | ENCSR185FOY |
| ZNF467 | HEK293 | Schmitges, et al. 2016<sup>82</sup> | GSM2026821 |
| YY1 | HEK293 | Schmitges, et al. 2016 | GSM2026776 |
| ZNF143 | HEK293 | Ngondo-Mbongo et al., 2013<sup>83</sup> | GSM959048 |
| EP300 | HEK293 | No publication | GSM1239071 |
| In situ Hi-C | Hela |  | 4DNFIE7V3DN9 |

## Data availability

All ChIP-seq, Micro-C, RNA-seq datasets generated in this study will be available from GEO upon request during the review process and released upon publication. Public datasets are provided through their accession codes from the related database.

## Code Availability

The related code for this study is available at https://github.com/boxialaboratory/3D-genome-ZJ.

## Acknowledgement

This work is supported by NIH grants DP5OD033430 and R03DE033371 (to B.X.). Arima Genomics partially supported this work through reagents. X.L. is a Fellow of The Jane Coffin Childs Fund for Medical Research. H.L. and B.X. are supported by the Junior Fellowship of the Harvard Society of Fellows. We thank Jason Buenrostro, Liz Gaskell, Fadi J. Najm, Noam Shoresh, Neva Cherniavsky Durand, and members of the Xia lab for valuable discussions and suggestions. We also thank the Genomics Platform and the Flow Cytometry Core at the Broad Institute for assistance with experiments. In addition, we thank Amelia Weber Hall, Guillermo Barreto Corona, Gwyneth Torrecampo, Bingxu Liu, Daniel Rosen, Kas Manakongtreecheep, Raktima Raychowdhury, Rollin Leavitt, and Amrita Vetticaden for sharing reagents and providing experimental assistance. Last, we are grateful to the ENCODE consortium for generating, organizing, and sharing many genomics datasets that were used in this study.

## Author Contribution

B.X. conceived the project. B.X. and J.B. designed the computational and experimental components and interpreted the results. J.T. and B.X. performed the in silico screen analyses with input from A.T. J.B. and Q.L. performed the genomic experiments with help from R.I., V.G., B.T., X.L. and A.S.H. J.B. performed the computational genomics analyses with help from J.T. and H.L. B.B. and A.S.H. provided feedback and contributed to discussion. B.X. supervised the study and led the writing of the manuscript with J.B. and Q.L. B.X. and J.B. edited the manuscript with key inputs from B.T. and A.S.H. All authors commented on the manuscript.

## Supplementary Figures

**Supplementary Figure 1.**
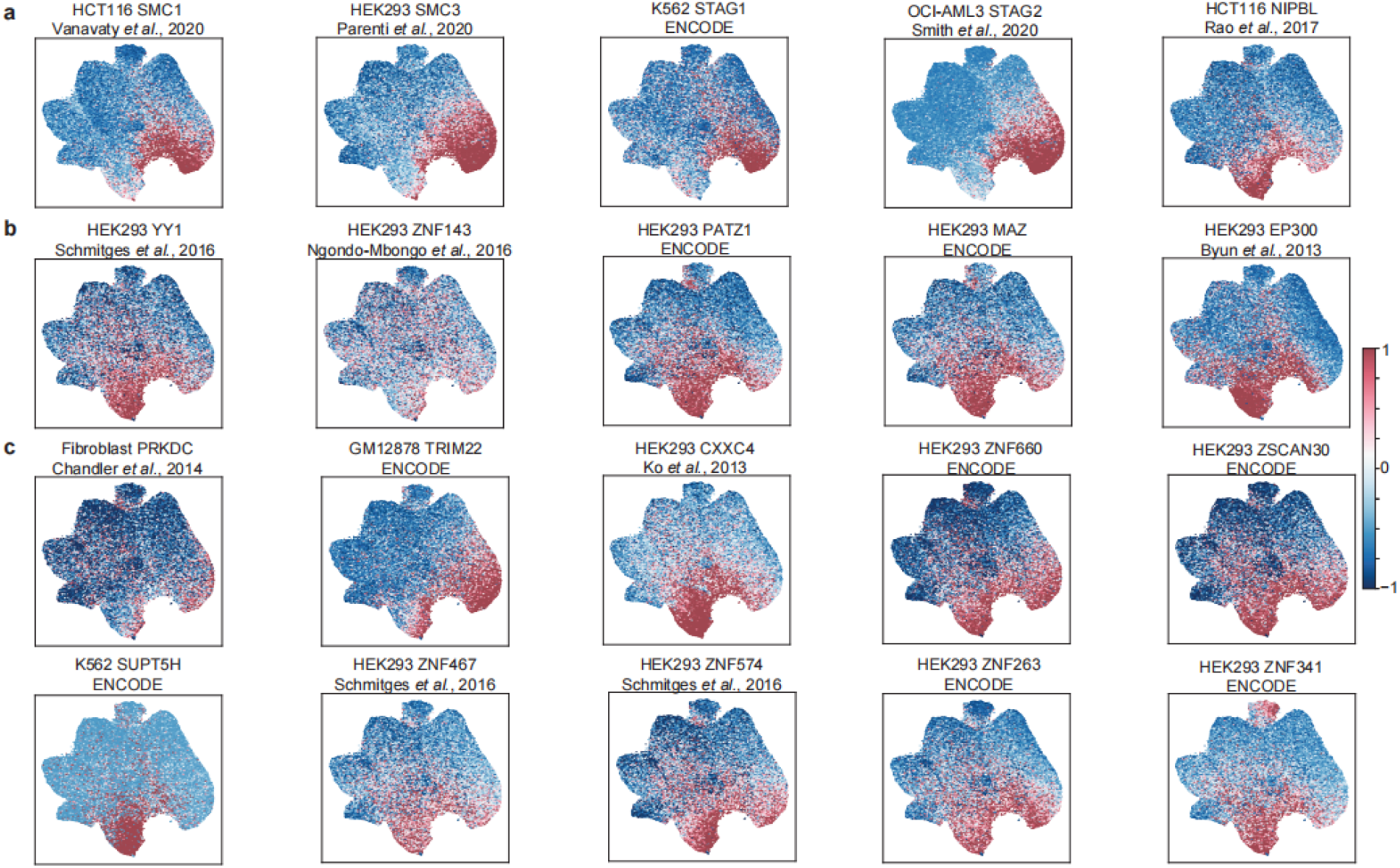
UMAP visualization of ChIP-seq signal intensities of additional candidate chromatin factors. **a**-**b**, Related to Fig. 1c, this UMAP embedding derived from HEK293 cells, and visualized chromatin factors for additional known 3D genome regulators (**a**), and recently proposed or putative regulators of chromatin interactions (**b**). Notably, these recently proposed or putative regulators of 3D genome organization tend to enrich at promoter-enhancer regions, with only minor overlap with CTCF-and cohesin-bound CRE cluster. **c**, Additional top-hit factors in the *in silico* screen (Fig. 1b).

**Supplementary Figure 2.**
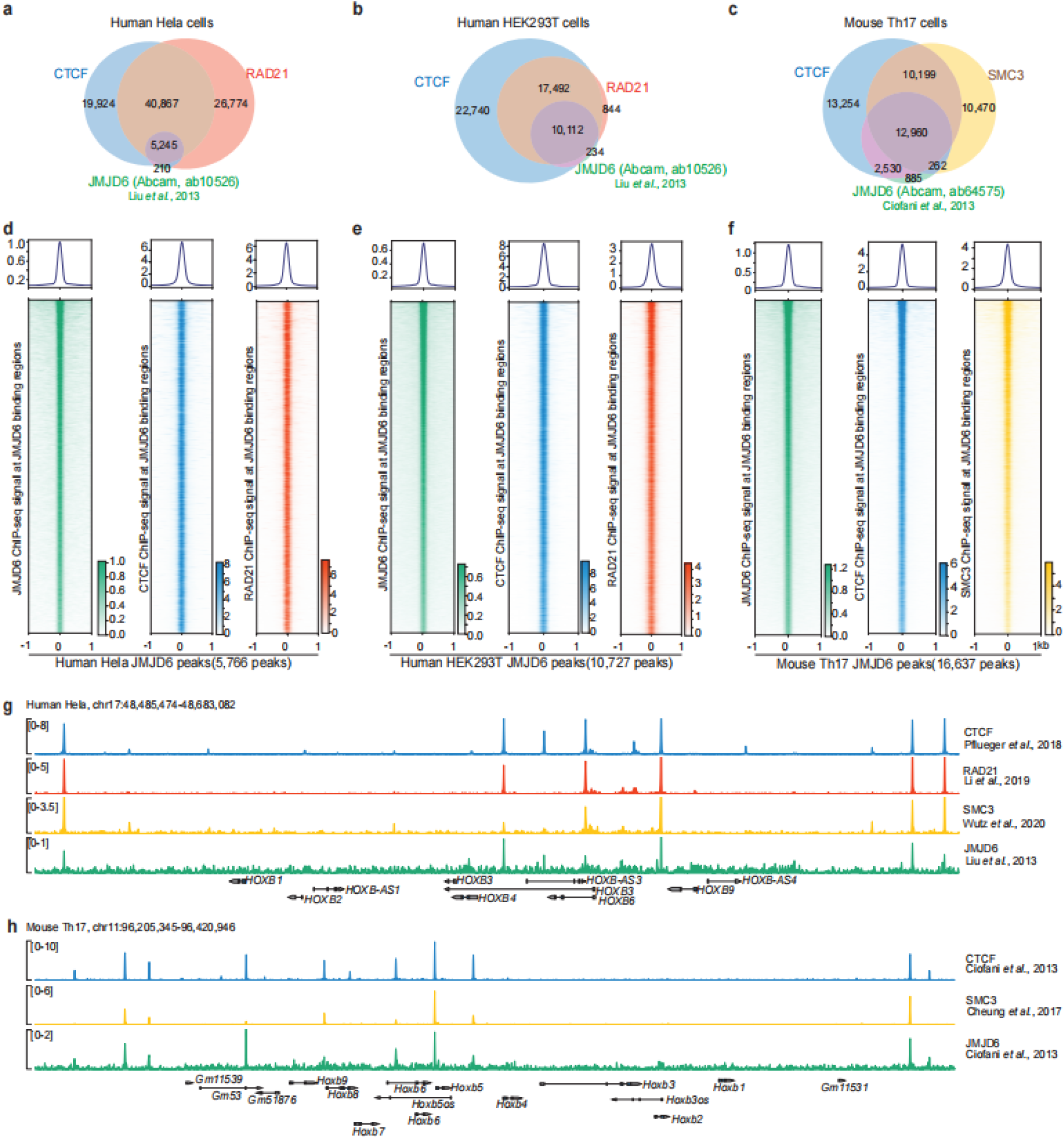
JMJD6 co-binds with CTCF, RAD21, and SMC3 across cell types and species. **a**-**c**, Overlap of JMJD6 binding sites with structural factors in (**a**) human HeLa cells (CTCF, RAD21), (**b**) human HEK293T cells (CTCF, RAD21), and (**c**) mouse Th17 cells (CTCF, SMC3). **d**-**f**, Heatmaps of ChIP-seq signals centered on JMJD6 peaks in HeLa (**d**), HEK293T (**e**), and mouse Th17 (**f**) cells. These results show that JMJD6 co-occupies CTCF and cohesin (RAD21 or SMC3) binding sites. Average binding intensities for each factor were shown above each heatmap. **g**-**h**, Genome browser views of JMJD6 and cohesin ChIP-seq signals at *HOXB* locus in human HeLa cells (**g**) and mouse Th17 cells (**h**).

**Supplementary Figure 3.**
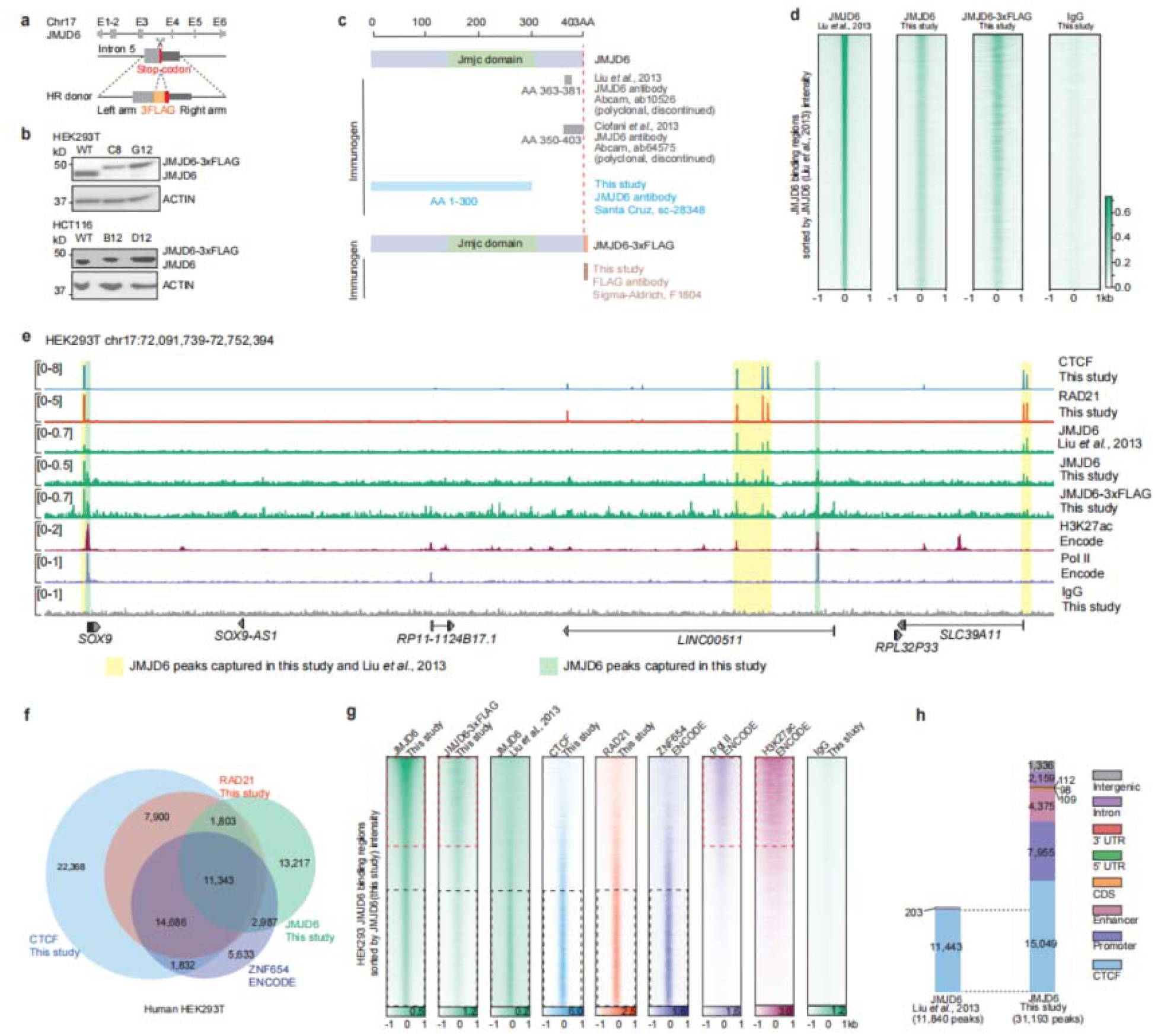
Validation of JMJD6 genome-wide binding via independent experiments. **a**, Schematic of CRISPR-mediated endogenous integration of 3×FLAG tag at the JMJD6 C-terminus by targeting the stop codon in exon 6. **b**, Western blot validation of JMJD6-3×FLAG protein in both HCT116 and HEK293T cells. **c**, Epitope positions of JMJD6 antibodies used in this study and prior publications, with antibody sources indicated. **d**, ChIP-seq signal heatmaps (±1 kb) centered on JMJD6 peaks from Liu *et al.* (2013), indicating that independent ChIP-seq experiments using a newly characterized JMJD6 antibody (N-terminus epitope) or anti-FLAG (for JMJD6-3×FLAG) recapitulate the JMJD6 binding sites from public data. **e**, Genome browser view of ChIP-seq tracks at the *SOX9* locus in HEK293T cells. Visualized ChIP-seq tracks include CTCF, RAD21, JMJD6 from public, JMJD6 from this study (independent anti-JMJD6 or anti-FLAG), H3K27ac, RNA Pol II, and IgG control. Yellow bars mark peaks shared between public JMJD6 ChIP-seq and the datasets from this study, whereas green bars highlight additional binding peaks at the active transcription-related chromatin regions discovered from independent JMJD6 ChIP-seq experiments (anti-JMJD6 or anti-FLAG). **f**, Venn diagram showing overlap among CTCF, RAD21, JMJD6 (anti-JMJD6 from this study), and ZNF654 (ENCODE) peaks in HEK293T cells. **g**, Heatmap visualization of all chromatin signals at the JMJD6 peaks identified in this study. Visualized signals include JMJD6, structural proteins (CTCF, RAD21, ZNF654), and transcription-associated chromatin signals (RNA Pol II, H3K27ac). Red dashed box (top) denotes regions enriched for transcription-associated signals, whereas black dashed box (bottom) highlights TAD boundary regions (strong binding by CTCF, ZNF654, and cohesin) while display minimal transcription-associated signals (RNA Pol II and H3K27ac). **h**, Genomic annotation of JMJD6 peaks detected from public JMJD6 ChIP-seq data (Liu *et al.*, 2013) and current datasets (anti-JMJD6).

**Supplementary Figure 4.**
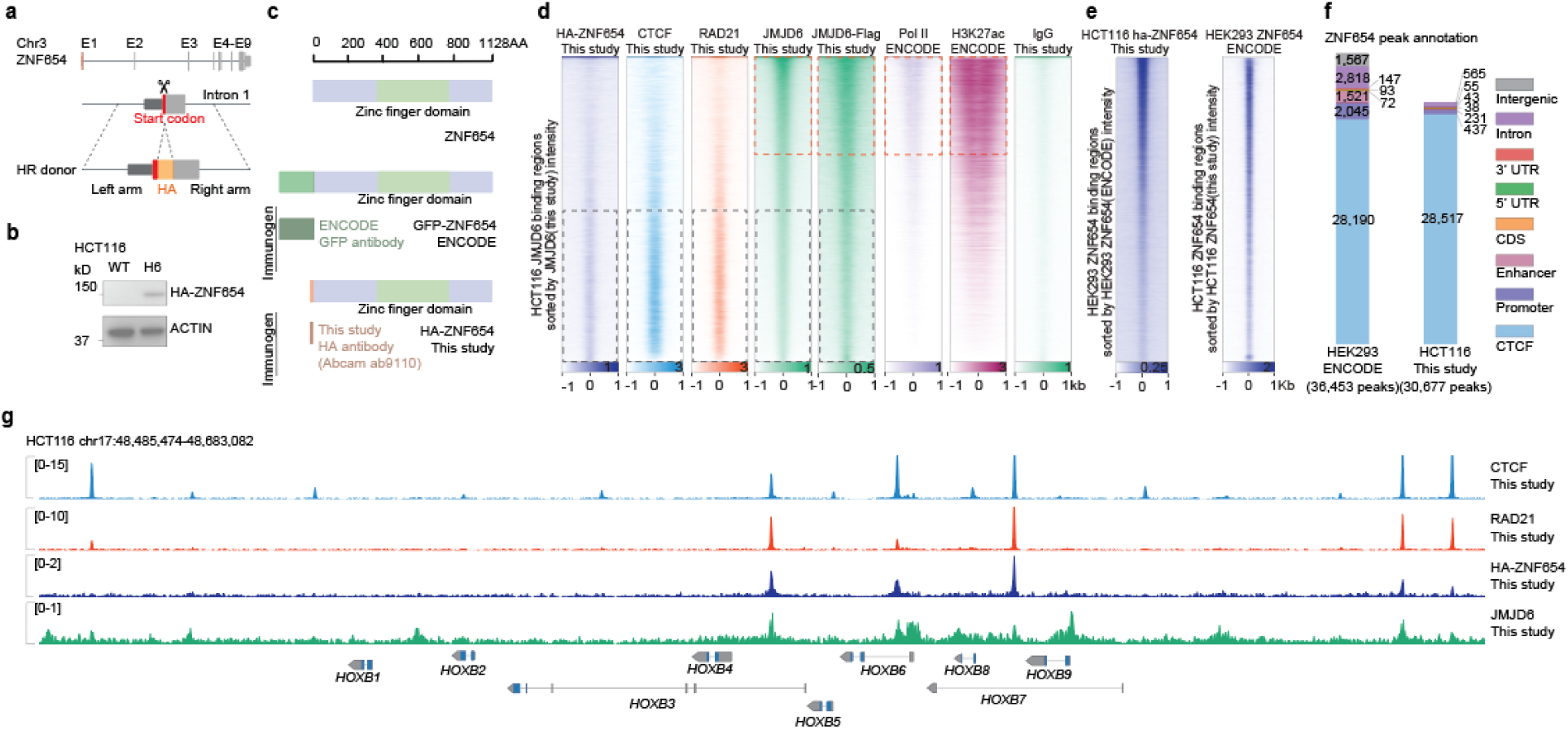
Validation of ZNF654 genome-wide binding via endogenous HA tagging. **a**, Schematic of CRISPR-mediated integration of an HA tag at the ZNF654 N-terminu by targeting the start codon in exon 1. **b**, Western blot validation of HA knock-in to ZNF654 in HCT116 cells. **c**, Protein domain structure of ZNF654 and the epitope tagging used in this study and in ENCODE datasets. **d**, ChIP-seq signal heatmaps (±1 kb) in HCT116 cells show ZNF654 co-localizes with CTCF and RAD21 but minimal overlap with active chromatin marks (H3K27ac, Pol II). Red dashed box (top) denotes regions enriched for transcription-associated signals, whereas black dashed box (bottom) highlights TAD boundary regions. **e**, Reciprocal heatmaps (±1 kb) of ZNF654 binding signals suggest its conserved binding across cell types. ZNF654 ChIP-seq in HCT116 cells (this study) plotted over HEK293 ZNF654 peaks (ENCODE), and vice versa. **f**, Genomic annotation of ZNF654 peaks detected from ENCODE dataset (in HEK293 cells) and from this study (anti-HA-ZNF654 in HCT116 cells). **g**, Genome browser view of the *HOXB* locus in HCT116 cells displaying ZNF654, JMJD6, CTCF, and RAD21 ChIP-seq tracks.

**Supplementary Figure 5.**
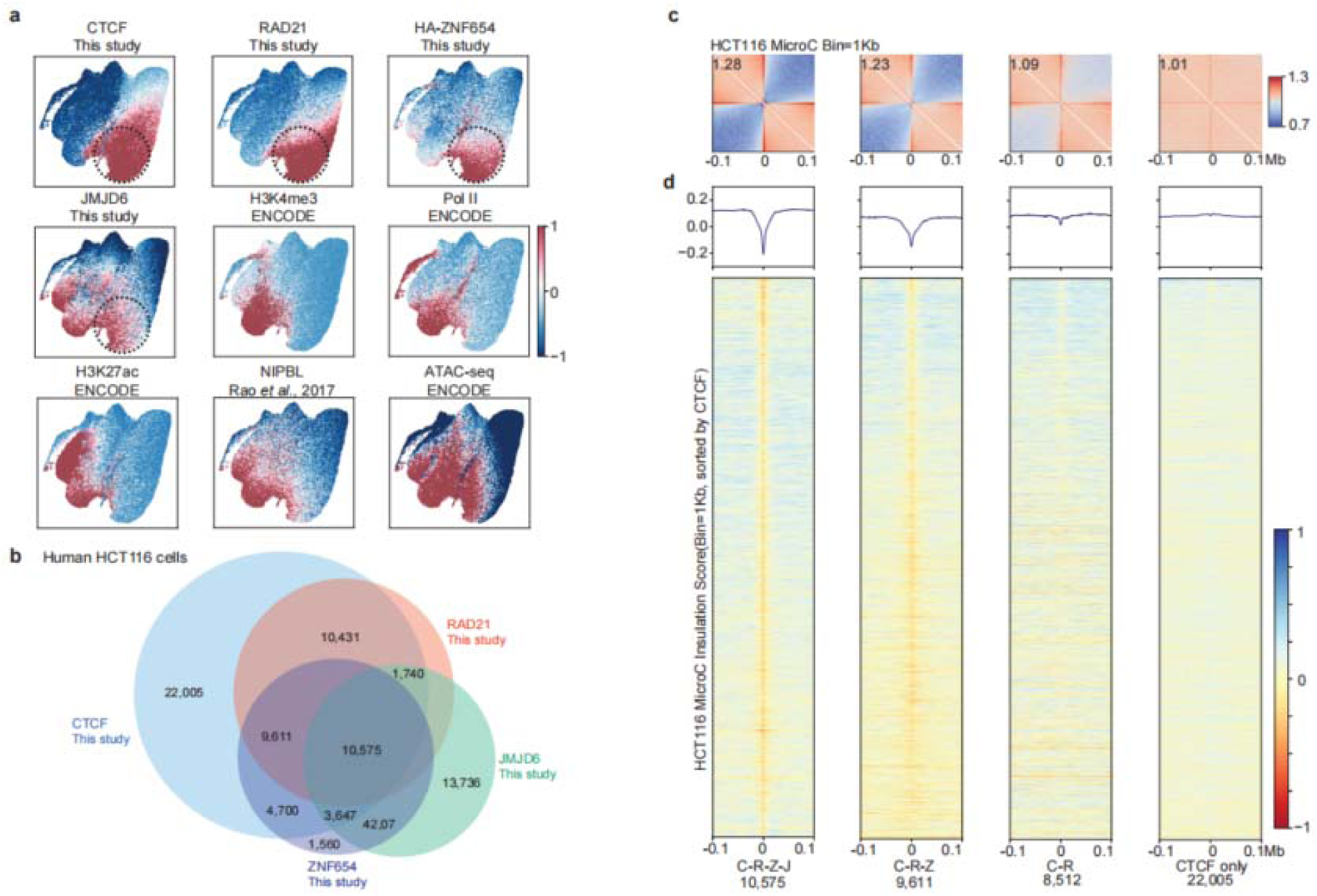
ZNF654 and JMJD6 uniquely define TAD boundaries in HCT116 cells. **a**, UMAP projection of chromatin feature ChIP-seq signal intensities across HCT116 cCREs UMAP embedding. The black circle marks the C–R–Z–J co-binding cluster, exhibiting a distribution highly consistent with that observed in HEK293 cells. **b**, Venn diagram showing peak overlaps among CTCF, RAD21, ZNF654, and JMJD6 in HCT116 cells. **c**, Micro-C pile-up visualization of chromatin interaction intensities flanking the loci belonging to the four CTCF-occupied groups in HCT116 cells. **d**, Heatmaps of insulation scores (at 5kb/bin resolution) centered on peaks from each binding category in HCT116 cells.

**Supplementary Figure 6.**
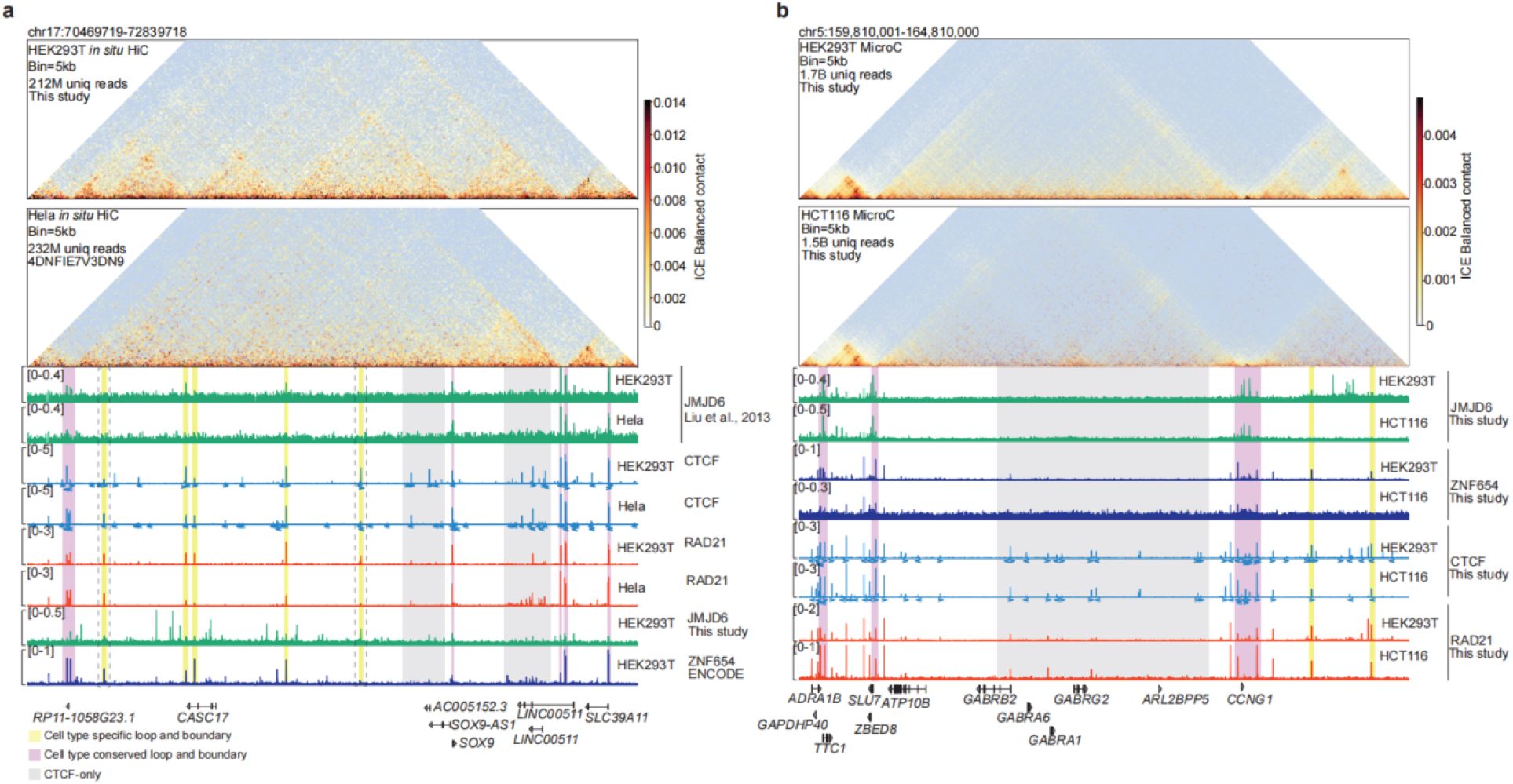
**Representative examples of cell type-specific TAD boundaries and chromatin loops**. **a**, *In situ* Hi-C contact maps (5-kb resolution) at the *SOX9* locus in HEK293T and HeLa cells, with corresponding JMJD6, CTCF, and RAD21 ChIP-seq tracks. Cell type-specific ZNF654/JMJD6 binding aligns with TAD boundaries in HEK293 cells, whereas CTCF and RAD21 binding is conserved in both cell types. The black dashed box highlights HEK293T-specific JMJD6 peaks detected with the new antibody but absent in public data, matching the HEK293T-specific boundary. **b**, Micro-C contact maps (5-kb resolution) at the *GABRA* gene cluster in HEK293T and HCT116 cells. Differences in domain organization correlate with ZNF654 and JMJD6 binding, despite conserved CTCF and RAD21 occupancy. Color shading indicates HEK293T-specific (yellow), conserved (purple), and CTCF-only (grey) loop and boundary regions.

**Supplementary Figure 7.**
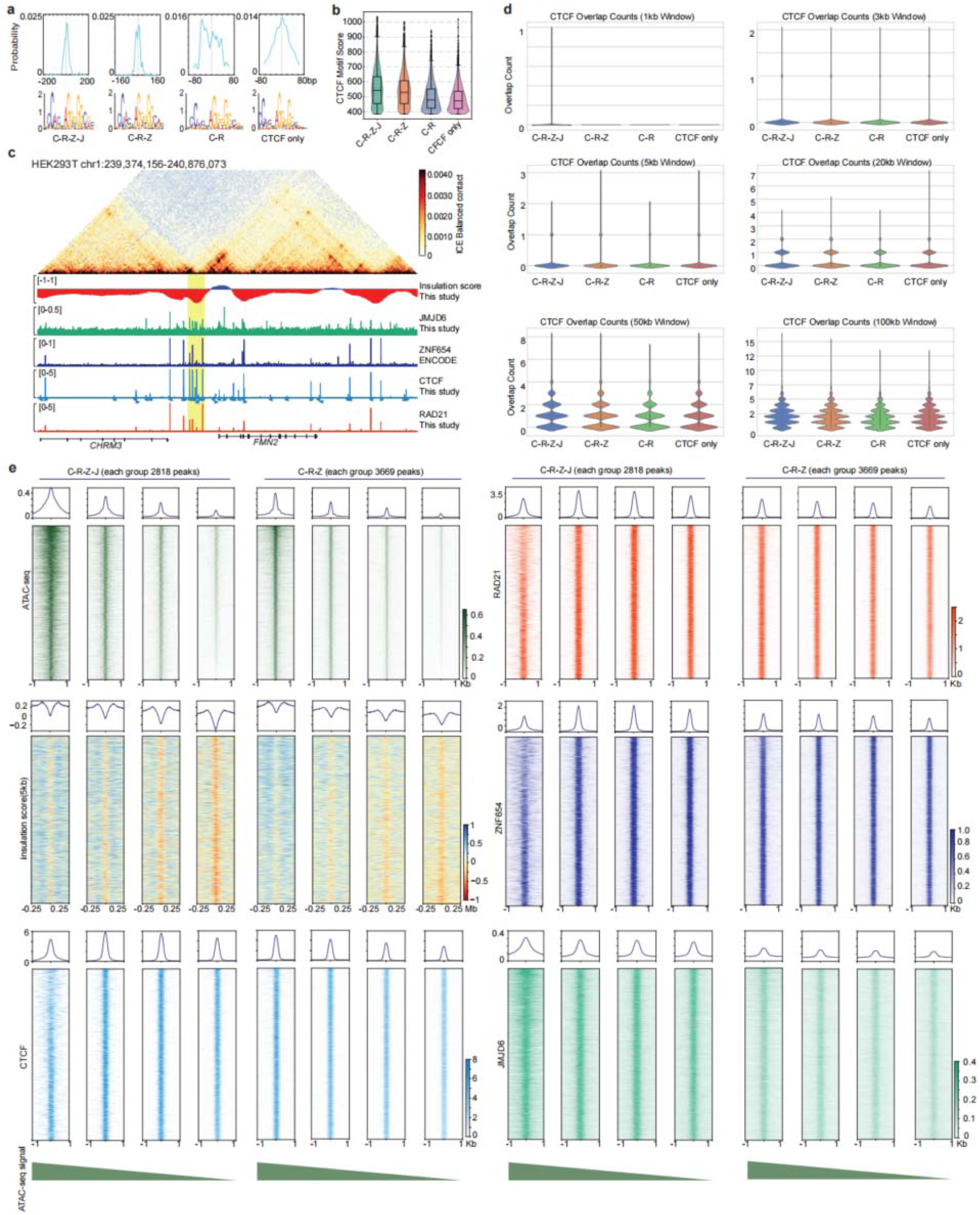
Interpreting sequence and chromatin feature difference between boundary and non-boundary CTCF sites. a, CTCF motif distribution and position frequency matrices (PFMs) across the four binding categories (C–R–Z–J, C–R–Z, C–R, and CTCF-only), showing similar motif patterns across groups. **b,** Violin plots of CTCF motif scores for each category, indicating no significant differences. **c,** Genome browser view of a representative HEK293T region showing a strong TAD boundary with multiple adjacent CTCF sites, such CTCF clusters contain peaks from all four binding categories. The Micro-C contact map resolution: 10kb/bin. **d,** Quantification of local CTCF site clustering within ±1 kb to ±100 kb windows across categories, showing no systematic differences. **e,** Heatmaps of ATAC-seq, CTCF, RAD21, ZNF654, JMJD6, and insulation score across four chromatin-accessibility quartiles within C–R–Z–J and C–R–Z peaks. Insulation strength increases as chromatin accessibility decreases, whereas CTCF, RAD21, ZNF654, and JMJD6 binding levels remain largely unchanged.

**Supplementary Figure 8.**
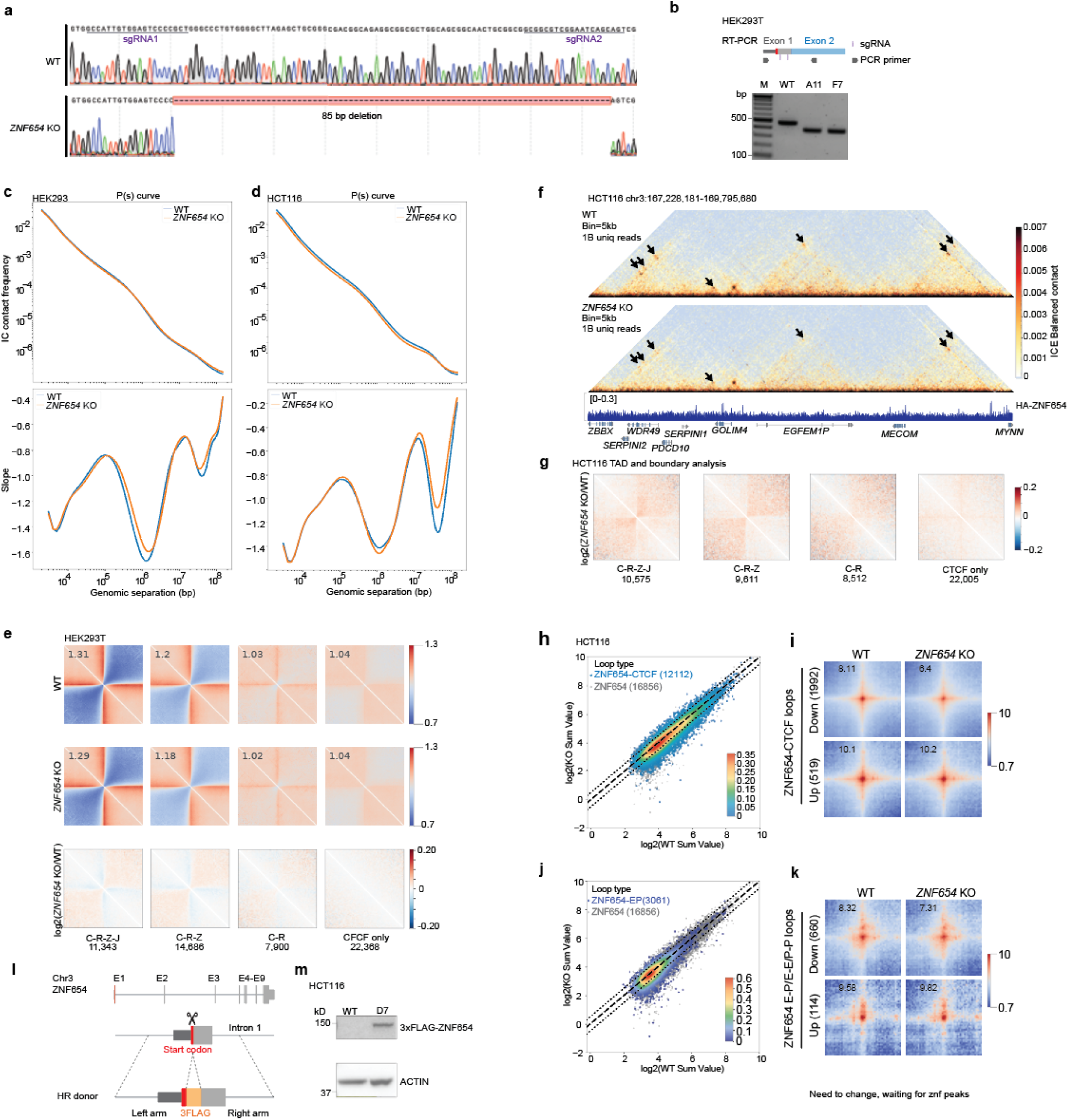
**ZNF654 strengthens TAD boundaries in both HEK293 and HCT116 cells**. **a-b,** Validation of *ZNF654* knockout in HEK293 cells by genomic PCR with Sanger sequencing (**a**) and RT–PCR (**b**). **c-d,** P(s) scaling curves and derived slopes for WT and *ZNF654* KO Micro-C libraries in HEK293T (**c**) and HCT116 (**d**), confirming comparable library quality. **e,** Pile-up Micro-C maps centered on the four binding categories in WT (top), *ZNF654* KO (middle), and difference (bottom) matrices in HEK293 cells, showing reduced boundary insulation upon knockout, particularly at C–R–Z–J and C–R–Z sites. **f,** Representative Micro-C maps at the *MECOM* locus in HCT116 showing weakened loops and boundaries in *ZNF654* KO cells. **g,** Pile-up analysis of interaction changes across CTCF-occupied groups, highlighting increased inter-TAD interactions at C–R–Z–J and C–R–Z loci following ZNF654 loss in HCT116. **h-i,** Scatter plot (**h**) and APA (**i**) comparing loop strength between WT and *ZNF654* KO at ZNF654–CTCF co-occupied loop sites in HCT116 cells. **j-k,** Same analyses as in (**h**-**i**) for ZNF654 occupied enhancer-promoter loops. **l–m,** Schematic (**l**) and validation (**m**) of N-terminal 3×FLAG–ZNF654 knock-in in HCT116 cells.

**Supplementary Figure 9.**
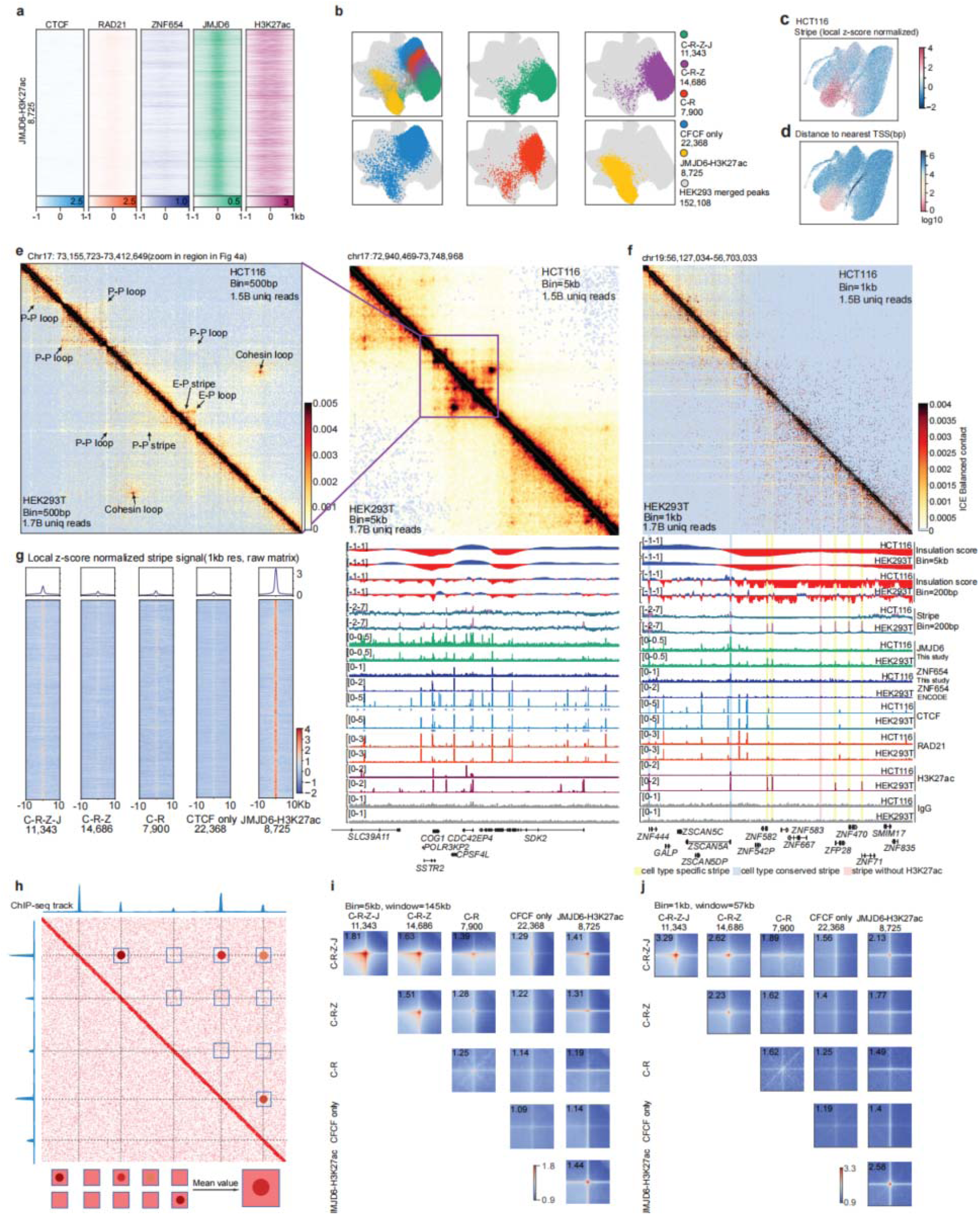
JMJD6 marks transcription-related stripes and enhancer–promoter/promoter–promoter loops. **a,** Heatmaps of ChIP–seq signal (±1 kb) centered on JMJD6–H3K27ac co-bound peaks in HEK293 cells for CTCF, RAD21, ZNF654, JMJD6, and H3K27ac. **b,** UMAP projection of factor binding categories (C–R–Z–J, C–R–Z, C–R, C-only, and JMJD6–H3K27ac) mapped onto the previously computed HEK293 UMAP space shown in Fig. 1c. **c,** Local z-score normalized stripe strength in HCT116 projected onto the same UMAP (shown in Fig S5a), showing enrichment near promoter regions. **d,** Distance to the nearest transcription start site (TSS) for each peak in HCT116 plotted in the same UMAP space. **e,** ICE-balanced Micro-C contact maps in HEK293T and HCT116 cells. At 5-kb resolution, a strong cohesin loop is observed (right, within purple box); a 1-kb resolution zoom-in reveals enhancer–promoter (E–P) and promoter–promoter (P–P) stripes that bypass the cohesion loop (left, arrow pointed cohesin loop. The zoom-in region, together with its corresponding ChIP–seq profiles and Micro-C stripe score, is shown in Fig. 4a. **f,** Comparison of cell-type specific stripes between HEK293T and HCT116, with JMJD6 marking stripe anchors more strongly than H3K27ac. **g,** Heatmaps of local z-score normalized stripe signal calculated from unbalanced matrix centered on five binding categories. **h,** Schematic of how ChIP–seq peaks are paired to form peak–pairs and how Micro-C contact frequency enrichment is calculated for each pair. **i-j,** Aggregate enrichment heatmaps of Micro-C contact frequencies centered on peak pairs at 5-kb (**i**) and 1-kb (**j**) resolution for the five categories (C–R–Z–J, C–R–Z, C–R, C-only, and JMJD6–H3K27ac), shown for all pairwise combinations.

**Supplementary Figure 10.**
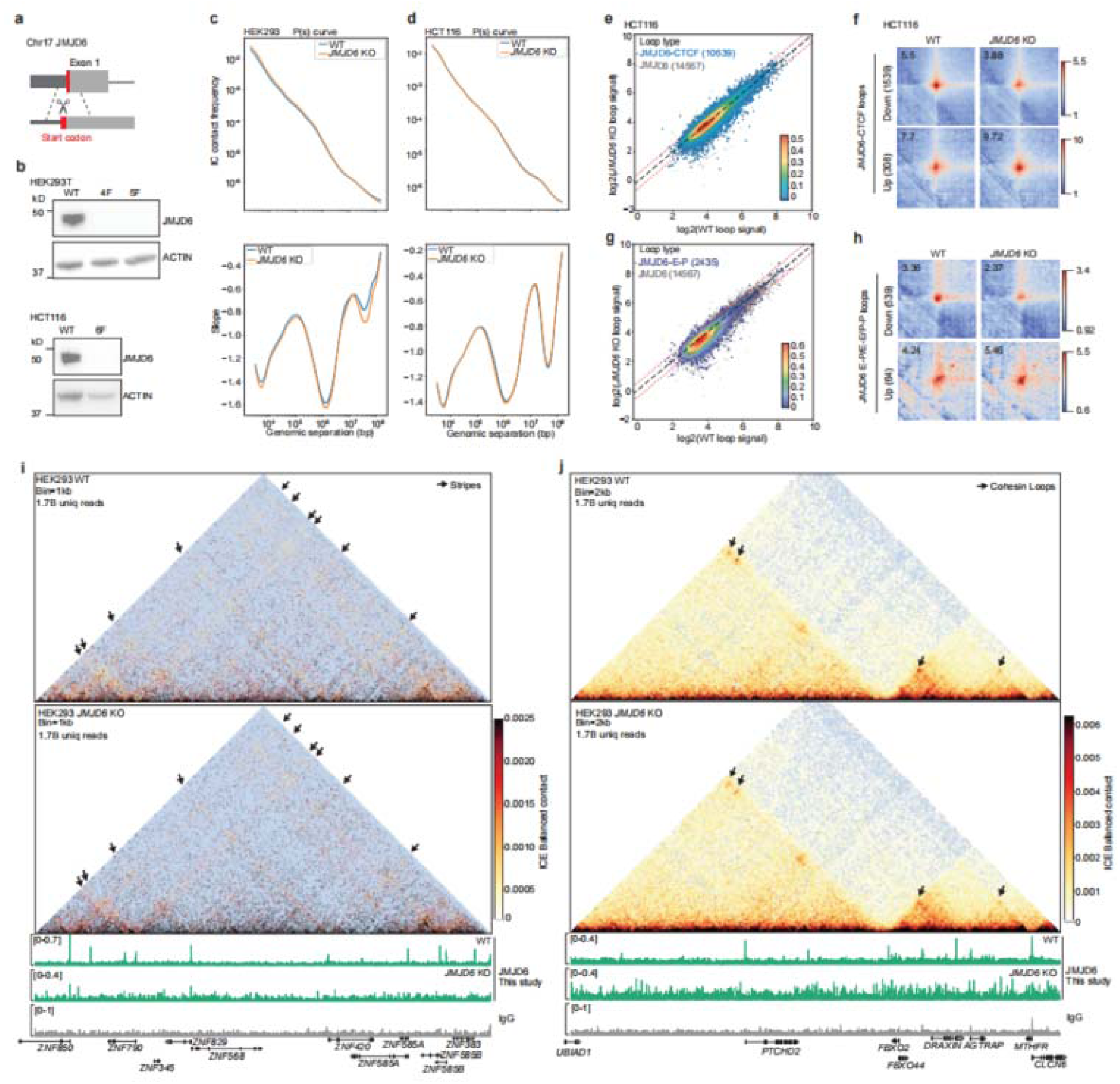
*JMJD6* knockout weakens cohesin loops and E-P/P-P loops and stripes. a, Schematic of CRISPR–Cas9 mediated *JMJD6* knockout. **b,** Western blot validation of *JMJD6* knockout in HEK293T and HCT116 cells. **c-d,** Contact probability scaling curves P(s) (top) and corresponding slopes (bottom) for WT and *JMJD6* KO Micro-C libraries in HEK293 (**c**) and HCT116 (**d**). **e,** Scatter plot comparing loop strength between WT and *JMJD6* KO at JMJD6–CTCF loop sites in HCT116 cells. **f,** Aggregate peak analysis (APA) of significantly weakened and strengthened JMJD6–CTCF loops in HCT116. **g,** Same analysis as (e) for JMJD6 associated enhancer–promoter (E–P) loops in HCT116. **h,** APA of significantly weakened and strengthened JMJD6 associated E–P loops in HCT116. **i,** Representative 1-kb resolution Micro-C contact maps in HEK293T WT and *JMJD6* KO cells showing reduced stripe strength upon JMJD6 loss. **j,** Representative 2-kb Micro-C contact maps in HEK293T WT and JMJD6-KO cells showing the effects of JMJD6 loss on cohesin loops.

**Supplementary Figure 11.**
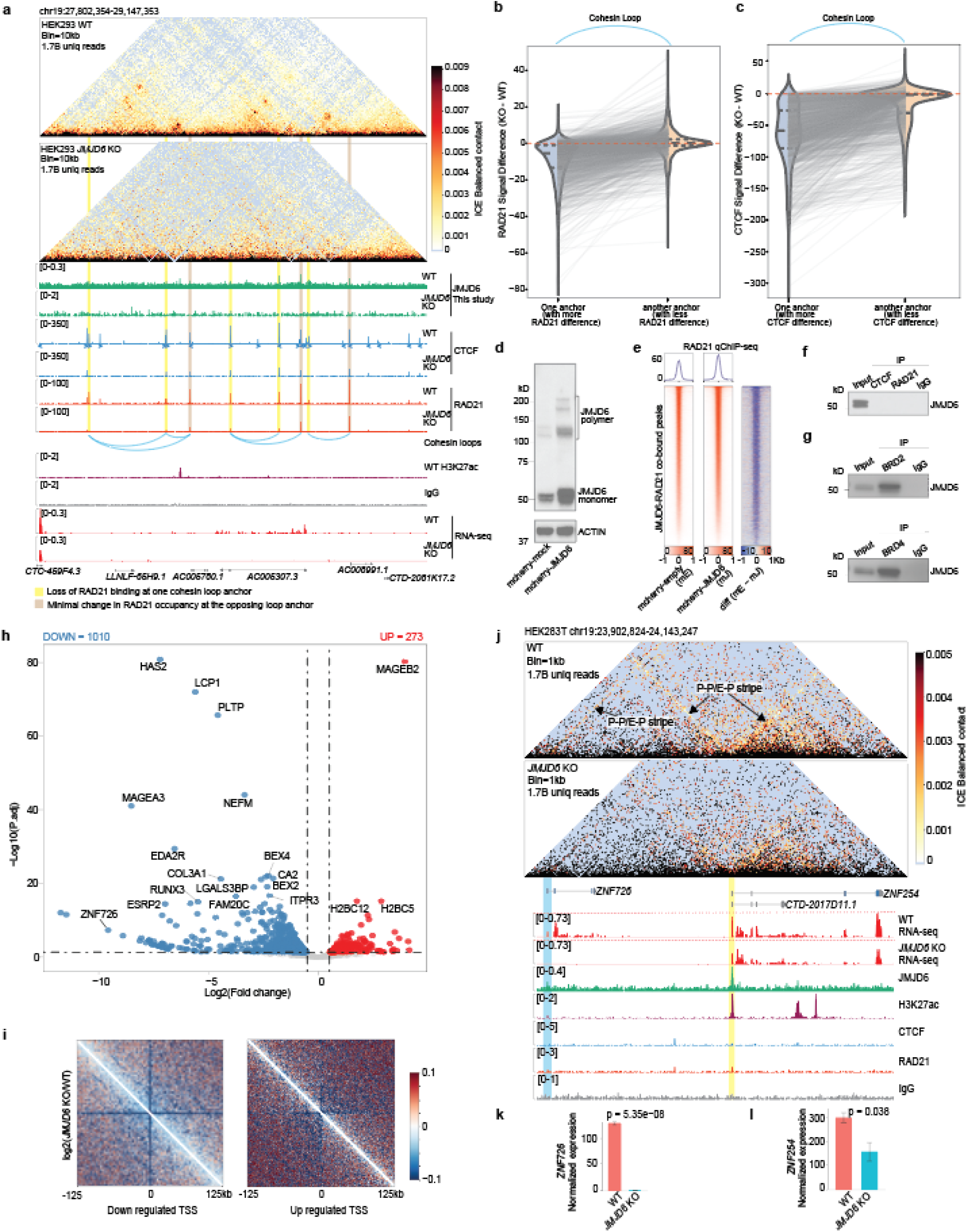
Loss of JMJD6 reduces cohesin occupancy and impairs promoter stripes and associated gene transcription. a, Representative Micro-C contact maps at the *AC005307.3* locus in HEK293T cells showing loss of cohesin loops upon *JMJD6* knockout. Loop loss is accompanied by reduced CTCF and RAD21 binding at one anchor of the loop. **b-c,** Genome-wide analysis of weakened cohesin loops, showing consistently reduced binding of RAD21 (b) and CTCF (c) at one loop anchor (anchors with more reduced occupancy are sorted to the left). **d,** Western blot validation of JMJD6 overexpression. **e,** RAD21 ChIP–seq signal increases at JMJD6–RAD21 co-bound sites upon JMJD6 overexpression. **f,** Co-immunoprecipitation shows that JMJD6 does not interact with CTCF or RAD21. **g,** Co-immunoprecipitation shows interaction between JMJD6 and BRD2/BRD4. **h,** Volcano plot of differentially expressed genes upon *JMJD6* knockout. **i,** Aggregate Micro-C maps centered on transcription start sites (TSSs) of downregulated and upregulated genes upon JMJD6 knockout. Downregulated genes show weakened promoter stripes, whereas upregulated genes display no stripe change but increased background interactions. **j,** Representative example showing that *JMJD6* knockout reduces *ZNF726* expression with complete loss of its promoter stripes, and reduces *ZNF254* expression with weakened promoter stripes. **k-l,** Bar plots showing significantly reduced expression of *ZNF726* (**k**) and *ZNF254* (**l**) upon *JMJD6* knockout.

---

**Supplementary Table 1.**
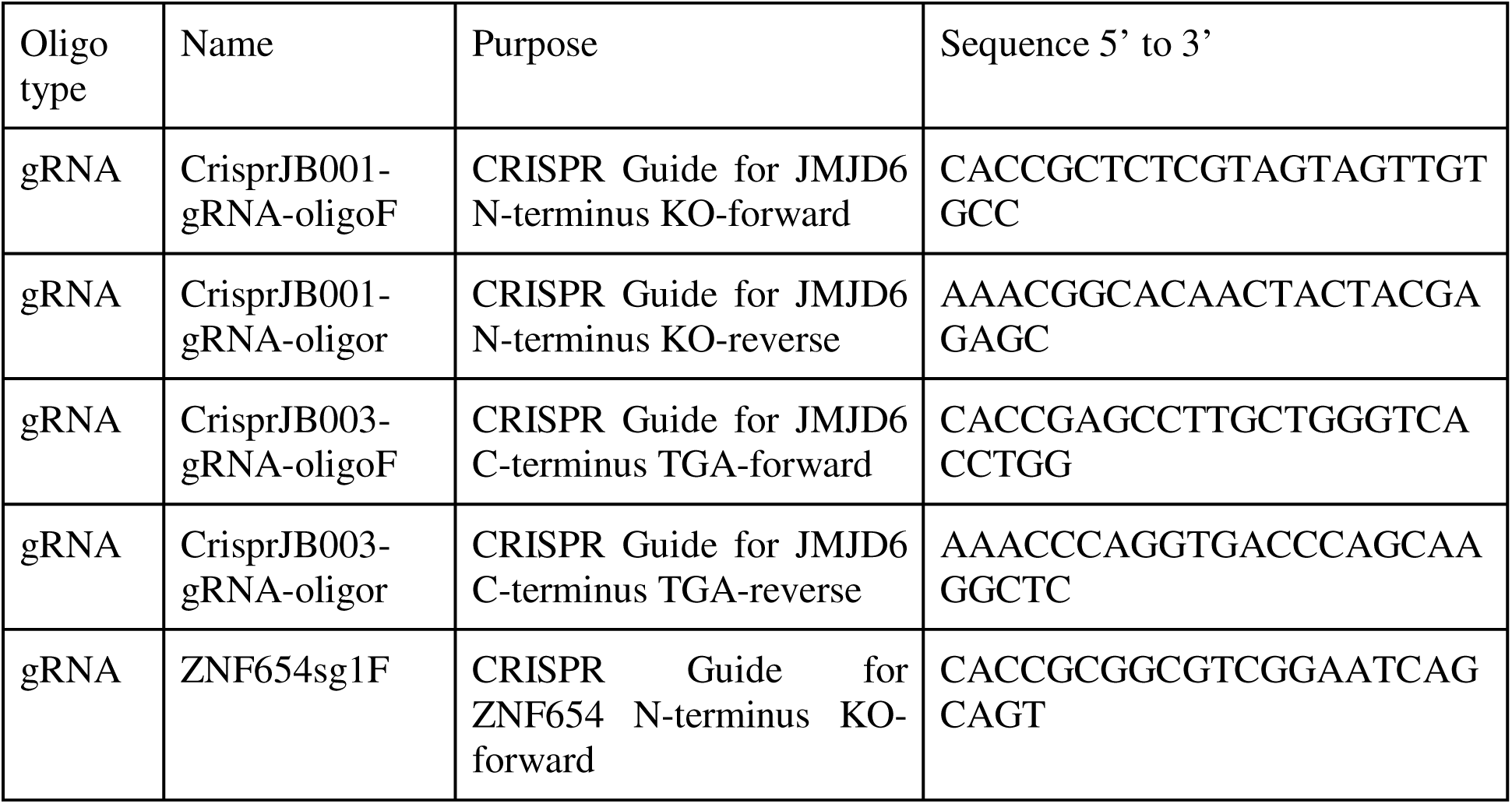
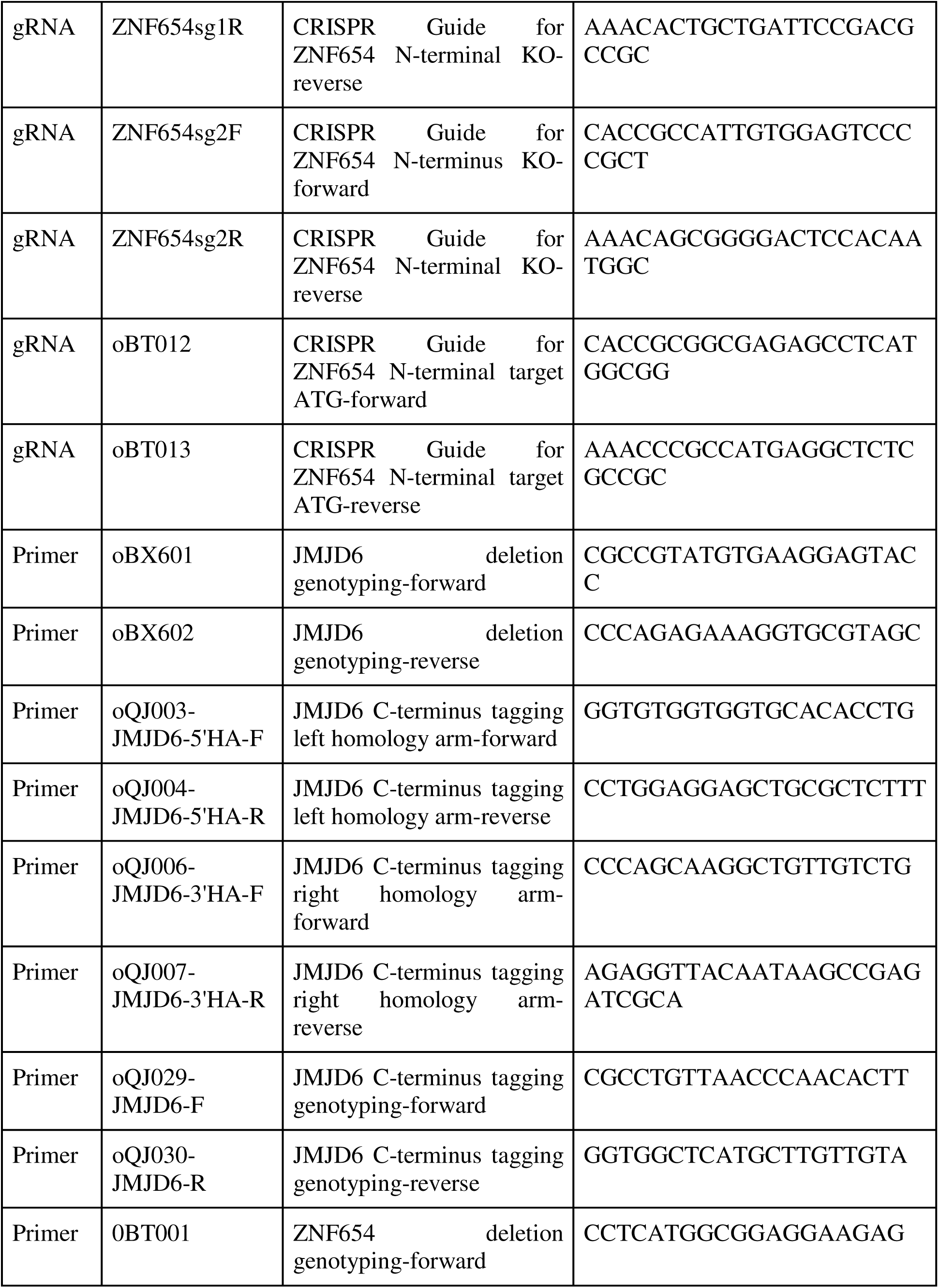
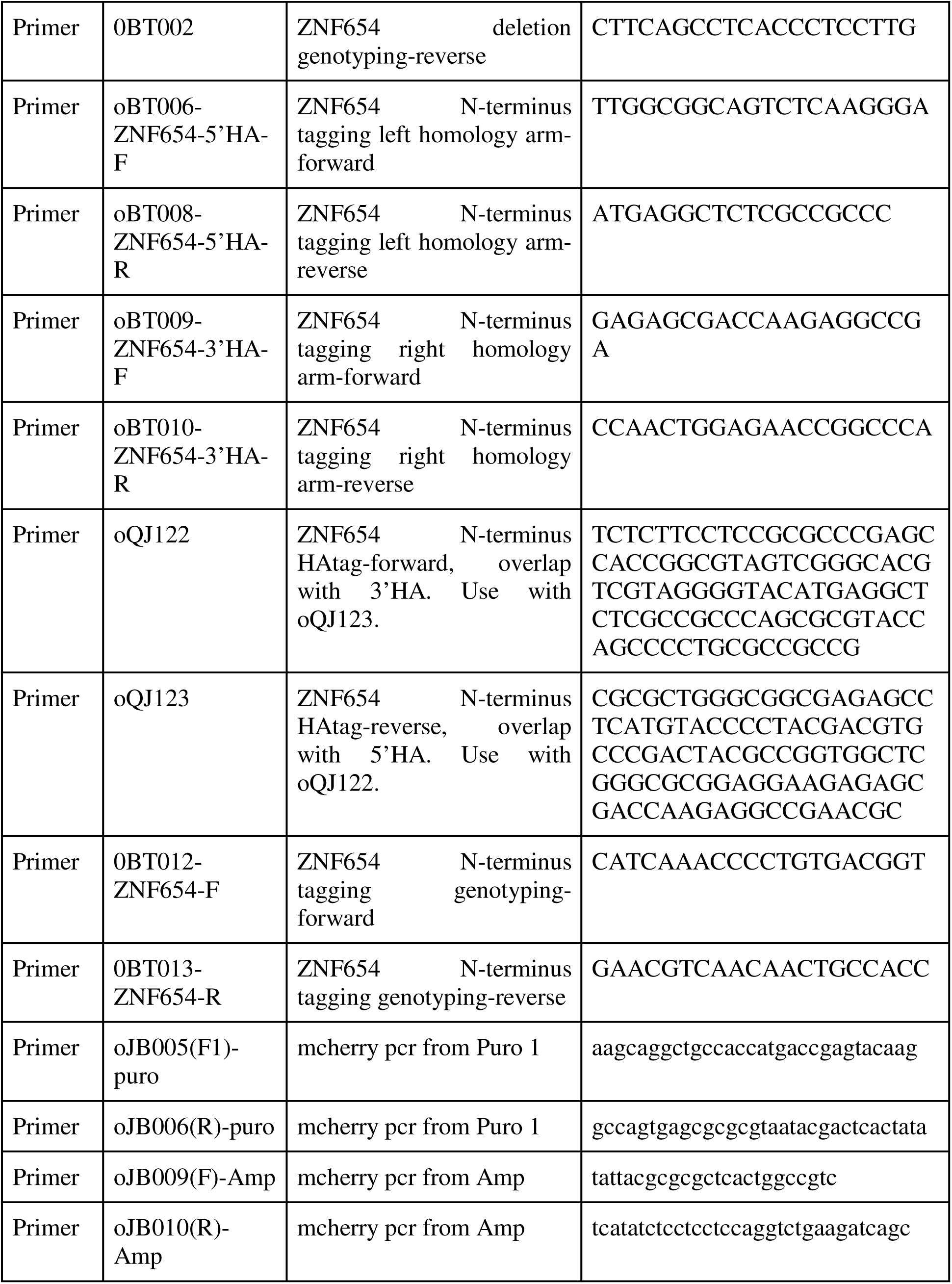
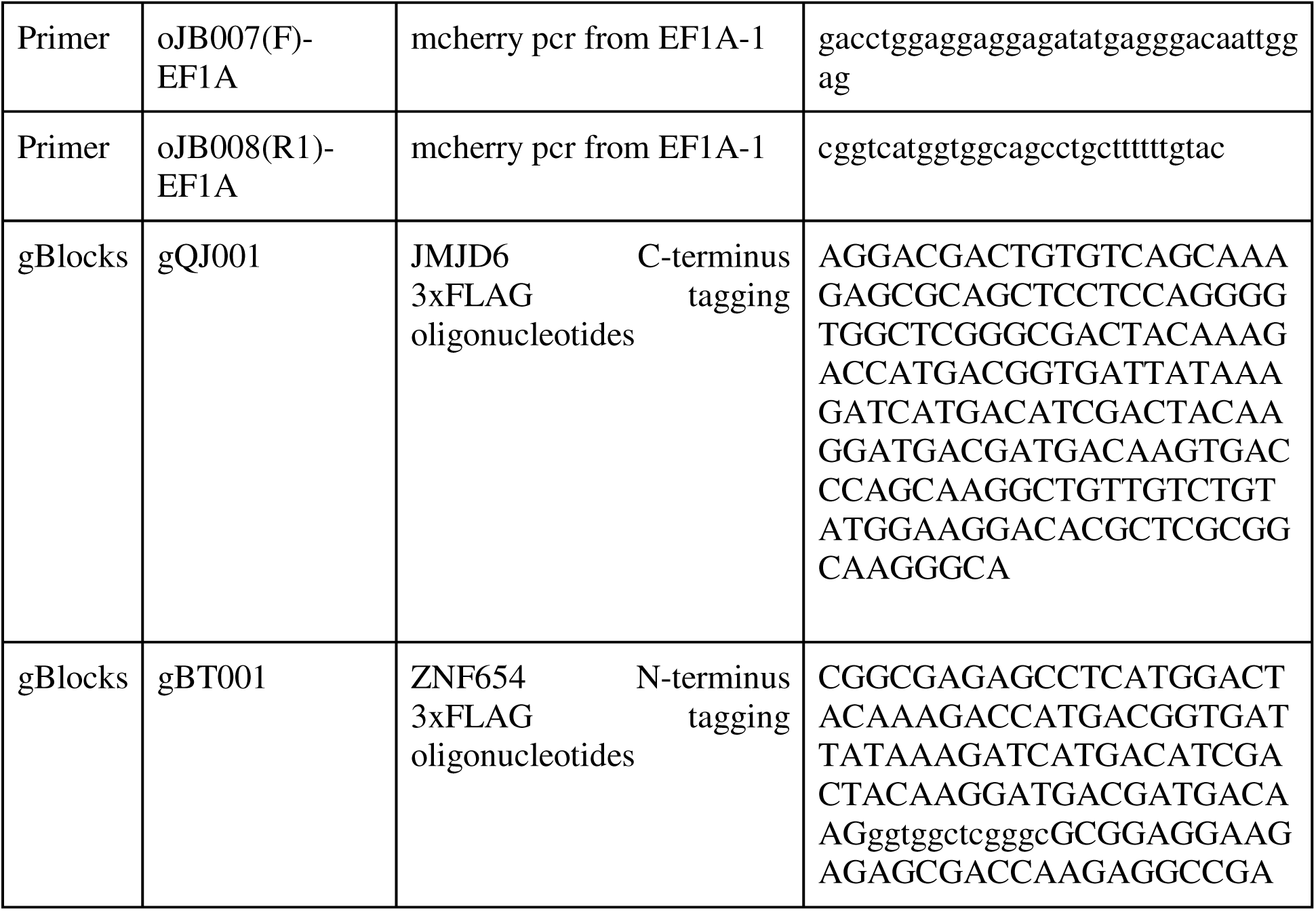

